# Top-down generation of low-resolution representations improves visual perception and imagination

**DOI:** 10.1101/2021.05.07.443208

**Authors:** Zedong Bi, Liang Tian

## Abstract

Perception or imagination requires top-down signals from high-level cortex to primary visual cortex (V1) to reconstruct or simulate the representations bottom-up stimulated by the seen images. Interestingly, top-down signals in V1 have lower spatial resolution than bottom-up representations. It is unclear why the brain uses low-resolution signals to reconstruct or simulate high-resolution representations. By modeling the top-down pathway of the visual system using the decoder of variational auto-encoder (VAE), we reveal that low-resolution top-down signals can better reconstruct or simulate the information contained in the sparse activities of V1 simple cells, which facilitates perception and imagination. This advantage of low-resolution generation is related to facilitating high-level cortex to form geometry-respecting representations observed in experiments. Moreover, our finding inspires a simple artificial-intelligence (AI) technique to significantly improve the generation quality and diversity of sketches, a style of drawings made of thin lines. Specifically, instead of directly using original sketches, we use blurred sketches to train VAE or GAN (generative adversarial network), and then infer the thin-line sketches from the VAE- or GAN-generated blurred sketches. Collectively, our work suggests that low-resolution top-down generation is a strategy the brain uses to improve visual perception and imagination, and advances sketch-generation AI techniques.

## Introduction

In perception, high-level cortex (such as prefrontal cortex) gets the information about the environment from low- level cortex (such as primary visual cortex, V1). It is believed that high-level cortex not only passively receives the feedforward input from low-level cortex, but also outputs signal to the low-level cortex through top-down pathway and adjusts its activity to minimize the difference between the top-down signal and the activity of low-level cortex [1]. This analysis-by-synthesis perspective of perception [2, 3] has received experimental support in both visual and auditory systems [3, 4]. Besides, when the subject is imagining a visual item, neural representations are found to be generated in a top-down cascade [5], and these representations are highly similar to those activated when the subject is looking at the item [6, 7]. These results above suggest that the brain performs like a top-down generator, which reconstructs or simulates the activity of the low-level cortex during perception or imagination.

However, in both perception and imagination, top-down representations have lower coding resolution than the bottom-up representations stimulated by seen images. During imagination, V1 neurons have larger receptive fields (RFs) and are tuned to lower spatial frequencies in imagined images than in seen images [8, 9]. During perception, the responses of V1 neurons to stimuli in classical RFs (mediated by bottom-up or feedforward connections [10]) are modulated by extra-classical RFs (mediated by top-down or feedback connections [11]) that surround classical RFs [12], which reflects the reconstruction of V1 activity by the higher-level cortex in in predictive coding formulations of perception [1]. These extra-classical RFs have larger sizes (i.e., lower spatial resolution) than classical RFs, and the modulation of neuronal activities by extra-classical RFs is insensitive to grating phase even for simple cells [13]. Why and how the brain uses the top-down low-resolution representations to simulate or reconstruct the bottom-up high- resolution representations in imagination and perception? Understanding the underlying computational principles is essential due to the fundamental role of perception and imagination in cognition [14, 15, 16, 17].

This paper addresses this problem by modeling the top-down pathway (from high-level brain area to V1) of the visual system as the decoder of variational auto-encoder (VAE) [18] (**Figure 1a**). VAE is a well-known image- generating paradigm, with the decoder transforming Gaussian distributed bottleneck states into meaningful images. In imagination or perception, the output of VAE decoder simulates or reconstructs the target bottom-up representation in V1 (**Figure 1b, c**). Interestingly, we find that for veridical imagination or perception, top-down signals should simulate or reconstruct the low-resolution representations (whose physiological candidates are V1 complex cells [19]), instead of the high-resolution representations of V1 simple cells (**Figure 1d**). This advantage of low-resolution top-down signals is related to the sparse activities of V1 simple cells [20] (**Figure 1d**). We further show that low-resolution top-down signals facilitate the high-level cortex to form representations that respect the geometric structure of the stimuli: two similar stimuli are represented by similar codes in the high-level cortex. Such geometry- respecting representations, which have been observed experimentally, are important for the behavioral flexibility (i.e., generalization in novel situations) of animals [21, 22, 23].

**Figure 1:**
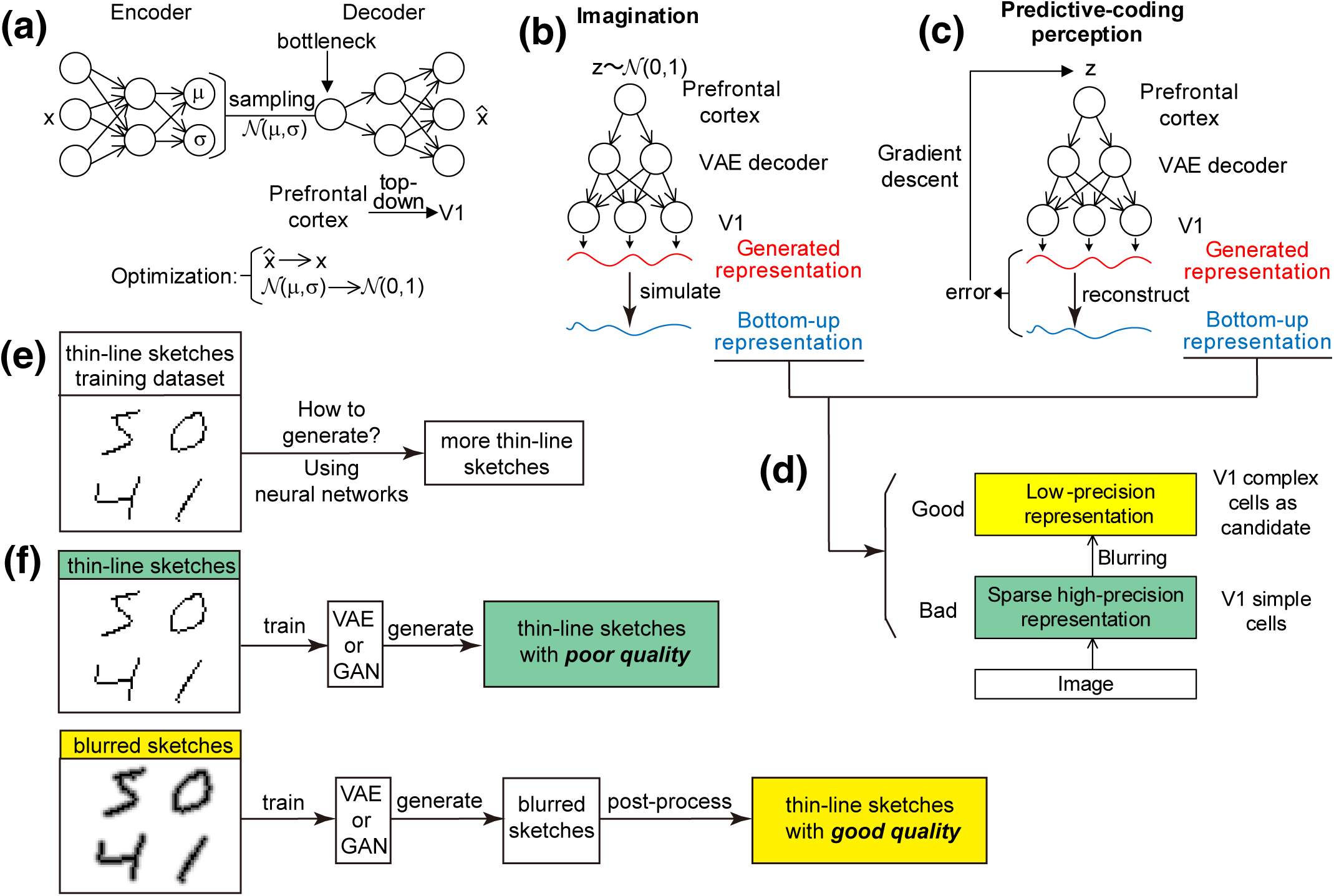
Model and main results of this paper. (**a**) Schematic of the variational auto-encoder (VAE). The value *z* of the bottleneck (which models the prefrontal cortex) is sampled from a Gaussian distribution *N* (*µ, σ*), with *µ* and *σ* determined by the two output channels of the encoder. This network is optimized so that the output *x̂* of the top-down pathway (decoder) is close to the input stimulus *x*, and at the same time *N* (*µ, σ*) is close to *N* (0, 1). After training, the encoder of VAE is abandoned, and the decoder is used to model the top-down pathway in the visual system in panels **b** and **c**. (**b**) Schematic of the imagination process. Only the decoder of a trained VAE is used. The bottleneck state *z* is sampled from the standard normal distribution, so that the generated images simulate target images in the real world. (**c**) Schematic of the predictive-coding perception process. The bottleneck state *z* is updated by gradient descent to minimize the error between the target image (blue) to be perceived and the generated image (red). (**d**) The bottom-up representations to be simulated or reconstructed by top-down signals during imagination or perception are not the representations of V1 simple cells, but the blurred representations of, possibly, complex cells. (**e**) The sketch-generation problem. Thin-line sketches are used to train neural networks to generate more sketches. (**f**) Upper panel: if VAE or GAN is trained directly using thin-line sketches, the generated sketches will have poor quality. Lower panel: we propose to train VAE or GAN using blurred sketches, and then post-process the VAE- or GAN-generated blurred sketches, resulting in good-quality generation of thin-line sketches.

Many studies have shown that understanding biological computation can boost artificial intelligence (AI) techniques [24]. Here we show that our above understanding of the biological visual system leads to a simple but effective AI technique to advance the generation of sketches (**Figure 1e**). Sketches are a style of drawings made of thin lines, which have been widely used in brain-storming creativity [25] and human-in-the-loop computer-aided artistic designing [26, 27, 28]. Thin-line sketches are more difficult to generate than ordinary images. With recent techniques, even a simple sketch requires multiple steps to generate, with each step generating a part of the sketch, in contrast with ordinary images that can be adequately generated in a single step [29]. It will be good if there is a technique that can generate high-quality sketches in a single step, thereby reducing the computational cost of sketch generation. In our technique (**Figure 1f, lower panel**), we first train VAE using blurred sketches, so that VAE can generate blurred sketches, corresponding to the low-resolution top-down signals in V1; then we infer thin-line sketches from the VAE-generated blurred sketches. This technique is different from traditional methods that train VAE using original thin-line sketches directly (**Figure 1f, upper panel**). We evaluated our technique using various datasets, such as the skeleton MNIST dataset [30], thin-line textures treated from OpenSimplex2 noises [31], and contours of CelebA faces [32], and found that our simple technique significantly improves the quality and diversity of the sketches generated in a single step. Besides, this technique is effective not only when we train VAE to generate sketches, but also when we train generative adversarial networks (GAN) [33] to generate sketches, demonstrating the broad applicability of the technique.

Collectively, our work explains the computation underlying the observations in biological experiments, and leads to an AI technique with broad applications.

## Results

### Model setup

We model the top-down pathway of the visual system using the decoder of a variational auto-encoder (VAE) [18] (**Figure 1a**), a popular neural network model for generating images. During training, the output of VAE is encouraged to reconstruct any given input, through a bottleneck state sampled from a Gaussian distribution whose mean and variance are determined by the output of the encoder part (**Figure 1a**), with this Gaussian distribution being encouraged to be close to the standard normal distribution. After training, the encoder of VAE is abandoned, and the decoder is used to model the top-down pathway of the visual system and is further used to study imagination and perception. The input of VAE decoder (i.e., the bottleneck of VAE) models the high-level cortex (such as the prefrontal cortex), and the output of VAE decoder models V1 (**Figure 1a**). Mental imagination requires top-down generation [6, 34], which can be modeled by sampling the bottleneck states from the standard normal distribution, so that the VAE decoder generates various outputs that simulate the bottom-up response of V1 to authentic images (**Figure 1b**). Predictive-coding perception [1] can be modeled by updating the bottleneck state to minimize the difference between the top-down generated image and the bottom-up response of V1 to the to-be-perceived image (**Figure 1c**). Therefore, VAE is an ideal generative model of the visual system, unifying imagination and perception.

There are two primary reasons that motivated us to choose VAE to model the visual system [15] (see more in the last paragraph of the Discussion):

(1) Our perception model is a predictive-coding model, in which the state of the high-level brain area is developed iteratively with the help of top-down generation (**Figure 1c**). In machine learning, compared with the perception using a pure feedforward network, this predictive-coding perception improves the robustness against adversarial attacks [35]. Biologically, it has been found that peripheral vision is more susceptible to illusions than foveal vision, which may be related to the lack of top-down connections to V1 areas responsible for peripheral vision [36, 37].
(2) The cost function that VAE minimizes is the variational free energy [18]. This is consistent with the free-energy principle of the brain [2], which posits that both perception and learning (and now imagination, too) can be understood by minimizing variational free energy. In machine learning, variational free energy is the negative evidence lower bound (ELBO) [38], which is the prediction accuracy minus the complexity of bottleneck-state representation. The assumption behind variational free energy minimization in our model is that the brain is trying to predict the world as accurately as possible while making a hypothesis about the world that is as simple as possible (an example of Occam’s principle). Compared with vallina autoencoders which focus on prediction accuracy without considering representation complexity, VAEs facilitate to develop smooth bottleneck-state representations that respect the shape morphing of images: in other words, there are no ’holes’ in the representation space of the bottleneck state that corresponds to unrealistic images at the output of VAE decoder, and two bottleneck states that are Euclideanly nearby also correspond to similar images at the output of VAE decoder [29, 39]. Such geometry-respecting representations are also observed in high-level brain areas such as prefrontal cortex and hippocampus [40, 41]. Developing this geometry-respecting representation is essential for semi-supervised learning [42, 43], few-shot learning [44], and data augmentation via imagination [44, 45].

Using VAE as a model of the visual system, we will study the computational advantage for low-resolution top-down generation (**Figure 1d**), and investigate the inspiration of this finding to advance the technique of sketch generation (**Figure 1e, f**).

### A computational model of V1 for imagination and perception

To validate the advantage of low-resolution top-down generation for imagination and perception (**Figure 1d**), we build a computational model of V1 and study the imagination and perception of the human faces in the CelebA dataset [32]. Below, we will first introduce how we model the activities of V1 simple cells and the corresponding low-resolution representation (**Figure 1d, red and yellow boxes**), and then illustrate our method to train the VAE and to study imagination and perception (**Figure 1b, c**).

To model the activities of V1 simple cells, we convolved the CelebA face images by Gabor filters with various spatial frequencies, orientations, and phases (**Figure 2b**), then passed the results through ReLU nonlinearity, obtaining many feature maps (**Figure 2a, middle column**). Each feature map was then treated by lateral winner-take-all (WTA). In lateral WTA, any pixel of a feature map (say, the red dot in **Figure 2c**) was compared with the nearby pixels beyond 30*^◦^* from the direction of the Gabor filter corresponding to the feature map (blue dots in **Figure 2c**). If the intensity of the red dot was larger than the intensities of the blue dots, the intensities of the blue dots were set to zero (see Methods). Lateral WTA resulted in thin lines in the feature maps, modeling the activities of simple cells (**Figure 2a, right column**). Lateral WTA models the lateral suppression between V1 simple cells [46, 47], which sharpens the tuning of V1 simple cells, especially under attention [48, 49]. This lateral suppression improves the sparseness of simple-cell activities [20], which reduces the representation redundancy along the direction perpendicular to the direction of the Gabor filter. There is no interaction between the red dot and the dots within 30*^◦^* from the direction of the Gabor filter in our model (grey dots within the circle in **Figure 2c**), which accommodates the contradicting experimental evidence for both collinear suppression and collinear facilitation in V1 [46, 50]. Overall, our model considers the heterogeneous recurrent interactions between V1 simple cells (i.e., the sign and strength of the recurrent connection between two V1 simple cells depend on the relative positions and orientation preferences of the two connected neurons) [51, 52], unlike the V1 model in [20] that regards simple cells as being independent with each other. However, unlike [51, 52], which integrated ordinary differential equations to model the dynamics emerging from these heterogeneous interactions, here we use lateral WTA to model this dynamics, which significantly simplifies the model and therefore reduces the computational cost to produce feature maps (**Figure 2a**) from CelebA images.

**Figure 2:**
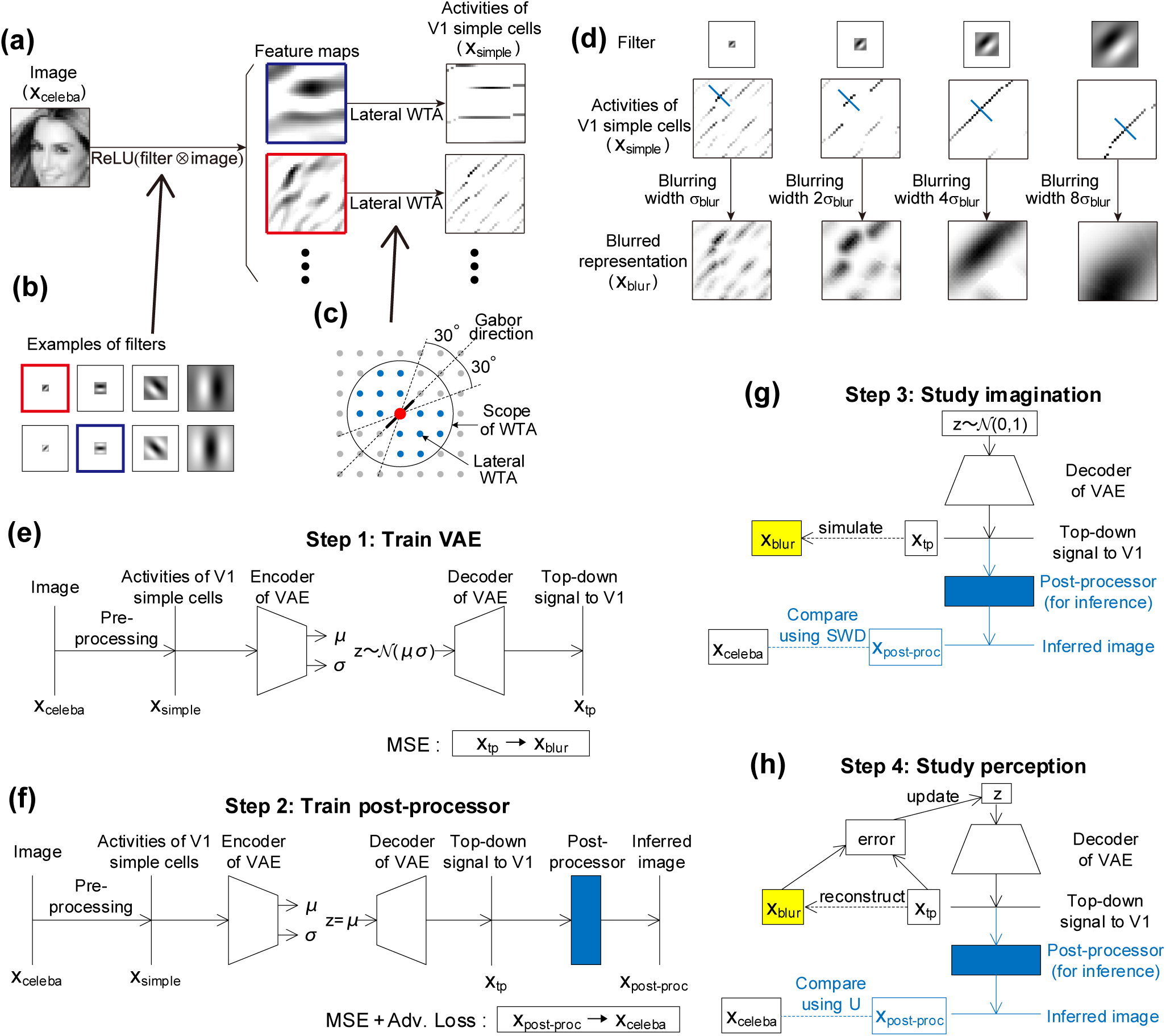
The computational model of V1 for imagination and perception. (**a**) The activities of V1 simple cells are modeled by convolving the image by Gabor filters, passing through ReLU nonlinearity, and then implementing lateral winner-take-all (WTA) to each feature map. (**b**) Examples of Gabor filters. The filters in the red and blue boxes respectively correspond to the feature maps in the red and blue boxes in panel **a**. (**c**) Schematic of lateral WTA. The thick bar indicates the direction of the Gabor filter in the red box of panel **b**. The neuron at the center (red dot) competes with the neurons indicated by the blue dots within the spatial scope of WTA (the circle) and beyond 30*^◦^* from the direction of the Gabor filter, so that if the red dot has the highest activity, the activities of the blue dots will be set to zero. The red dot does not compete with the grey dots. (**d**) To model the low-resolution representation (the yellow box of Figure 1d), we blur the activities of simple cells along the direction (blue bar) perpendicular to the direction of the Gabor filter. The blurring width corresponding to the Gabor filter with the highest spatial frequency is *σ*_blur_ (leftmost column), and is proportionally larger for a Gabor filter with lower spatial frequency (the other columns). (**e-h**) Our method to train VAE and study imagination and perception. (**e**) We inputted the simple-cells activities *x*_simple_ pre-processed from CelebA images *x*_celeba_ (panel **a**) to the VAE encoder, and trained the top-down signal *x*_tp_ to approach the blurred representation *x*_blur_ (panel **d**). Notice that the bottleneck state *z* is sampled from *N* (*µ, σ*) during training. (**f**) To train the post-processor (blue box), we let the bottleneck value of the VAE be the *µ*-channel of the encoder, and let the post-processed image *x*_post-proc_ approach the input image *x*_celeba_ by minimizing MSE loss plus adversarial loss (see Methods) . VAE is kept unchanged during the training of the post-processor. (**g**) To study imagination, we sample the bottleneck state from *N* (0, 1) and compare the inferred images treated by the post-processor with *x*_celeba_ using SWD (see Methods). (**h**) To study perception, we iteratively minimize the difference between the generated images by the VAE decoder and the blurred representation (yellow box) of the to-be-perceived image, and compare the inferred image with the to-be-perceived image using *U* (see Methods).

Imagination or perception requires top-down signals to simulate or reconstruct some target representation in V1 (**Figure 1b, c**). To model this target representation, we blurred the activities of simple cells along the direction perpendicular to the direction of the Gabor filter (**Figure 2d, blur bars**), using the parameter *σ*_blur_ to control the blurring width (**Figure 2d**).

We then trained a neural network model to illustrate the computational advantage of low-resolution top-down generation. We first trained a VAE with the activities of simple cells (**Figure 2a, right column**) being the input of the VAE and the blurred representations (**Figure 2d, bottom row**) being the target of the VAE output (**Figure 2e**). Then we trained another feedforward neural network (with U-Net structure, see the **blue box in Figure 2f**) as a post-processor to infer face-looking images (see **Figure 3a, b** as examples) from the blurred-line representations (**Figure 2d, bottom row**) generated by the VAE decoder: in this way, the information of the face-looking images contained in the blurred-line representations generated by VAE can be inferred, and used to quantify the quality of imagination and perception by comparing with the face images in the CelebA dataset (blue dashed lines in **Figure 2g, h**). By adjusting the parameter *σ*_blur_ (thereby controlling the spatial resolution of the blurred representations to train the decoder output of VAE, see **Figure 2d**) and studying the resulting quality of imagination and perception, we can draw a conclusion about how the spatial resolution of top-down signal to V1 may influence the quality of imagination and perception.

**Figure 3:**
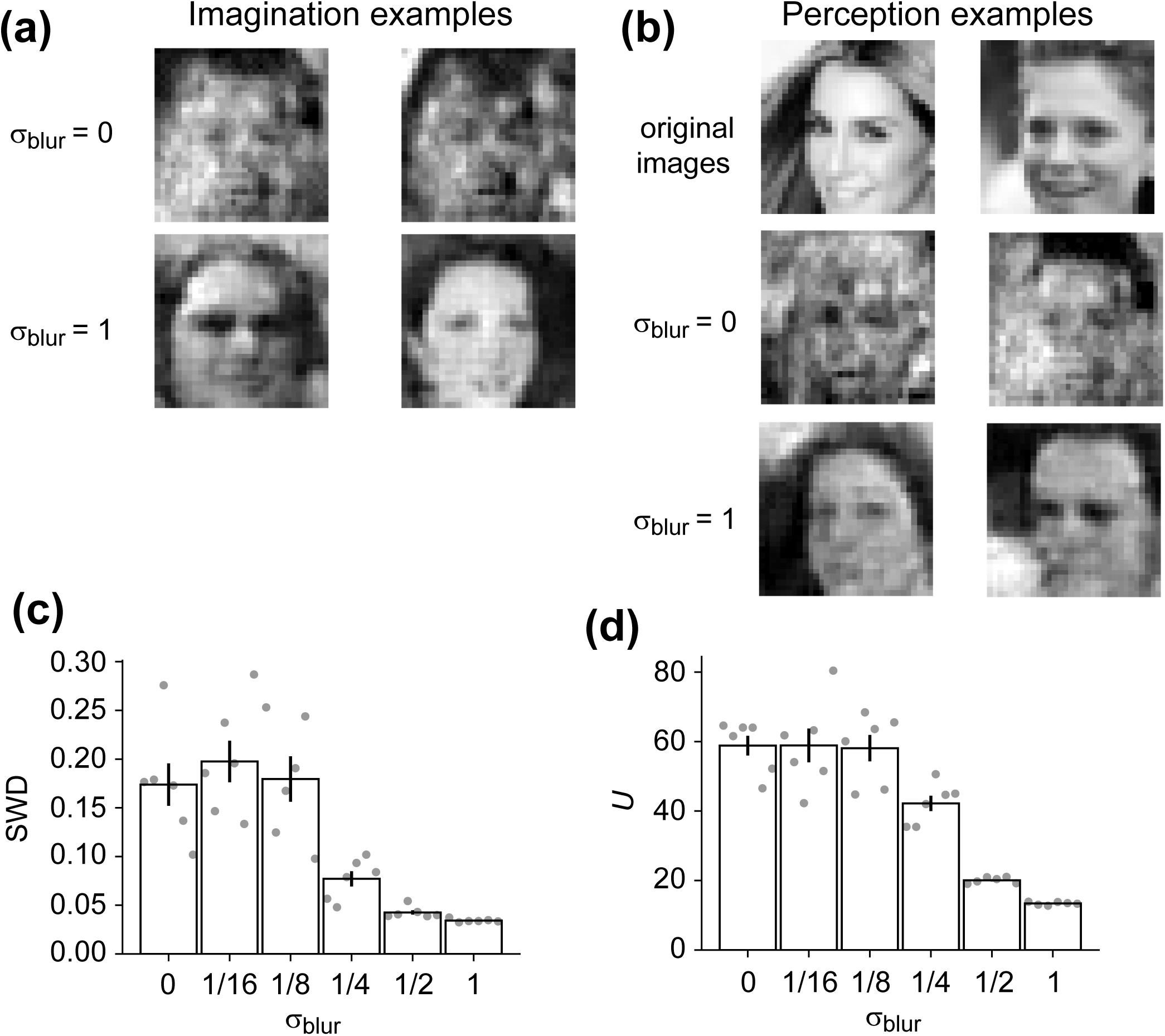
Low-resolution top-down generation facilitates imagination and perception. (**a**) Examples of the post-processed generated images in imagination when *σ*_blur_ = 0 or *σ*_blur_ = 1. (**b**) Examples of the post-processed reconstructed images in perception when *σ*_blur_ = 0 or *σ*_blur_ = 1. (**c**) Sliced Wasserstein distance (SWD) between the post-processed generated images in imagination and the images in CelebA dataset. (**d**) Difference *U* between the post-processed reconstructed images in perception and the to-be-perceived images in CelebA dataset. In panels **c** and **d**, each dot represents the result of a single VAE and post-processor configuration. Error bars represent s.e.m. over 6 configurations.

Imagination quality was evaluated by the sliced Wasserstein distance (SWD) [53, 54] between the CelebA images and the inferred images by the post-processor from the output of VAE (**Figure 2g**). In the calculation of SWD, we first projected the two datasets (i.e., CelebA images and the inferred images) in the same high-dimensional space onto the same one-dimensional space (i.e., a line), and then calculated the difference between the two projection distributions on the line. SWD is defined as the average of this difference over several random projection directions (see Methods). Perception quality was evaluated by the difference *U* between the CelebA images and the inferred images, after the difference between the output of VAE decoder and the target representation is iteratively minimized (**Figure 2h**). By adjusting *σ*_blur_, we could adjust the coding resolution of the target representation during training and, therefore, the output of VAE after training (**Figure 2d**). With the increase of *σ*_blur_, SWD decreases (**Figure 3c**), which means that the imagined images are more like the real images (**Figure 3a**); *U* also decreases (**Figure 3d**), which means that the top-down generated images are more like the to-be-perceived images (**Figure 3b**). In other words, low-resolution top-down signals facilitate imagination and perception.

Notice that the treatment from thin lines to blurry lines in **Figure 2d** corresponds to the computation performed by the connections from simple cells to complex cells in V1 [19]; the post-processor (**blue box in Figure 2f**), however, has no biological correspondence, but is merely our method to evaluate the quality of top-down generation.

### A simple but effective technique of sketch generation

We now study the AI technique to generate thin-line sketches (**Figure 1e**).

In the model above, the activities of V1 simple cells are thin lines in each feature map (**Figure 2a, right column**). When these thin lines are inputted to VAE, the best information recovery by the output is realized when the output generates blurred lines (**Figure 3**). This immediately implies a simple AI technique to generate sketches (drawings made of thin lines [28]) using VAE: instead of directly training VAE to generate thin-line sketches (**Figure 1f, upper panel**), we train VAE to generate blurred sketches, and then infer the thin lines from the generated blurred lines (**Figure 1f, lower panel** and **Figure 4a**). Here we investigate the advantage of this technique.

**Figure 4:**
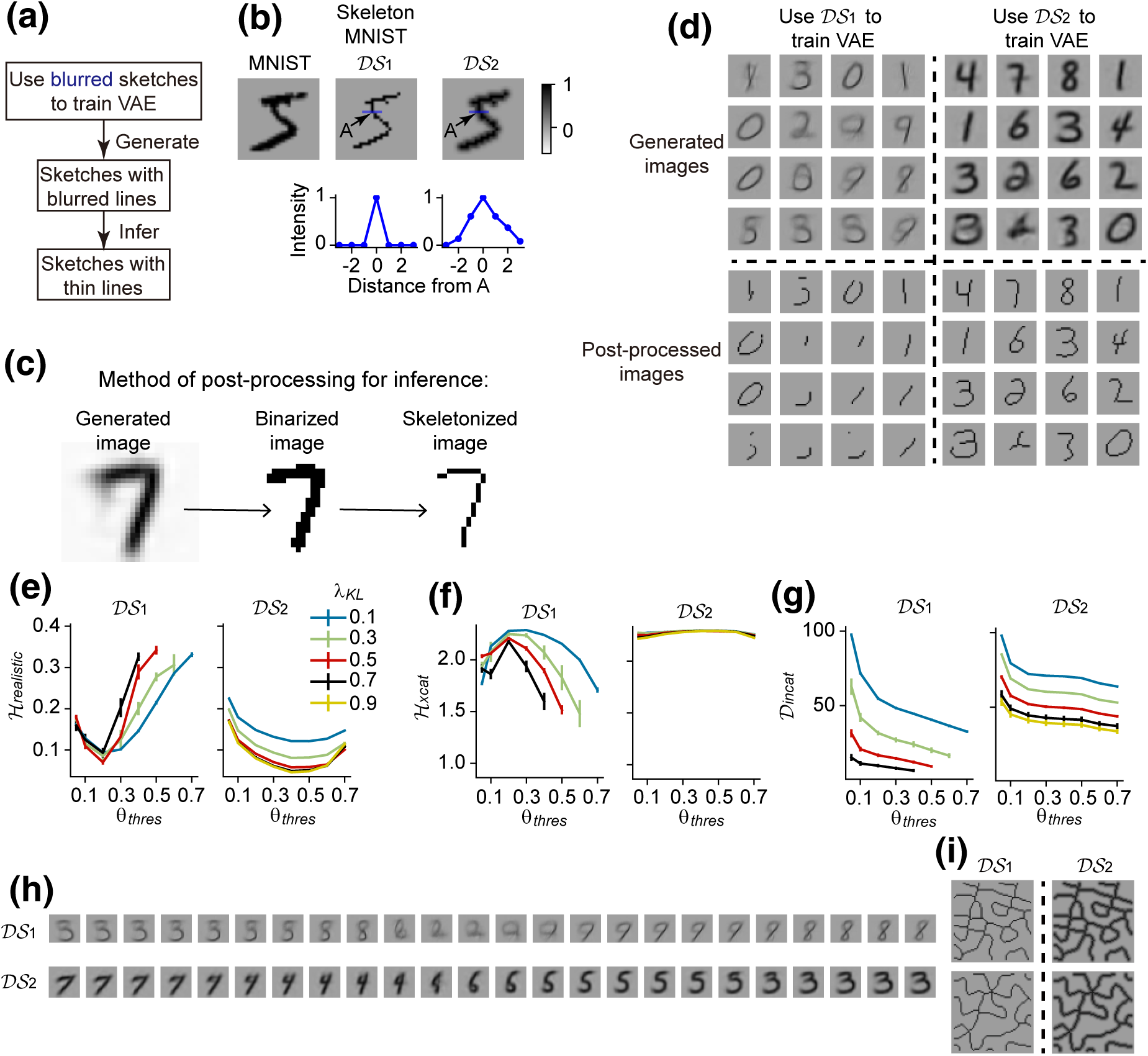
Low-resolution generation facilitates the generation of skeleton MNIST digits. (a) The proposed technique to generate sketches made of thin lines. (**b**) Upper subplots: example images in the MNIST, skeleton MNIST (*DS*_1_), and *DS*_2_ datasets. Lower subplots: pixel intensity along the blue bar in the upper panel as a function of the distance from point *A* (i.e., the middle point of the bar); note that it delta peaks for *DS*_1_ but decays slowly for *DS*_2_. (**c**) An example illustrating the post-processing of the VAE-generated images. (**d**) Upper subplots: examples of the generated images when using datasets *DS*_1_ (left subplots) and *DS*_2_ (right subplots) to train VAE. Lower subplots: the generated images after post-processing. Note that *DS*_2_ results in better image quality than *DS*_1_. (**e**) *H_realistic_* as a function of the binarization threshold *θ_thres_* when the parameter *λ_KL_* in VAE (see Methods) takes different values, when VAE is trained using *DS*_1_ and *DS*_2_ respectively. Some data points for *DS*_1_ are missing, see Methods for explanation. (**f, g**) Similar to panel **e**, but for *H_xcat_* and *D_incat_*. (**h**) Examples of the generated images when the bottleneck state slowly and linearly changes its value, see more examples in Figure S2. (**i**) Examples of the textures of thin lines that we also examined. We found similar results to MNIST digits (see Figure S5). In panels **e, f, g**, error bars representing s.e.m. over 8 VAE configurations. In panel **d**, *θ_thres_* = 0.4. In panels **d** and **h**, *λ_KL_* = 0.7.

We first conducted our study using the skeleton MNIST dataset [30]. Images in this dataset (denoted as dataset *DS*_1_ below) represent digits using lines of 1-pixel width (**Figure 4b, second column**). To study the best way to generate these 1-pixel-width lines, we trained VAEs using two datasets *DS*_1_ and *DS*_2_, with *DS*_2_ blurred from *DS*_1_ (**Figure 4b**).

To infer the 1-pixel-width lines from the VAE-generated images, we post-processed the VAE-generated images in the following way: we first binarized the images using a threshold *θ_thres_*, such that if the intensity of a pixel was larger (or smaller) than *θ_thres_*, the intensity would be set to be 1 (or 0), then we skeletonized the images (see Methods), resulting in binary images with lines of 1-pixel width (**Figure 4c and lower subplots of d**). By comparing the quality of the post-processed images resulting from the two datasets (*DS*_1_ and *DS*_2_), we can know the best way to generate thin-line sketches.

We quantify the quality of the generated digital images from three aspects. (1) Realisticity (quantified by *H_real_* ): a generated image should look like a digit in dataset *DS*_1_. (2) Cross-category variety (quantified by *H_xcat_*): the numbers of the generated images looking like different digits should be almost the same. In other words, it is not good if the images generated by a VAE all look like the same digit. (3) Within-category variety (quantified by *D_incat_*): the shapes of the images of the same digit should be various. Realistic and variable generation of digital images are manifested by small *H_real_*, large *H_xcat_* and large *D_incat_* values. *H_real_* indicates the quality of individual images, while *H_xcat_* and *D_incat_* indicate the diversity of images. See Methods for details.

We investigated *H_real_*, *H_xcat_* and *D_incat_* with the change of the binarization threshold *θ_thres_* and a parameter *λ_KL_* that controls the regularization strength onto the distribution of the bottleneck state variable in VAE (see Methods) . We found that VAEs trained by *DS*_2_ generated images with better eye-looking quality than VAEs trained by *DS*_1_ (**Figure 4d**), consistent with the quantitive indications of smaller *H_real_*, larger *H_xcat_* and larger *D_incat_* of the post-processed images in a large range of parameters *θ_thres_* and *λ_KL_* (**Figure 4e-g**, Figure S3). These results suggest that generating low-resolution representations improves the quality of the generated thin-line sketches.

To gain insight into why training VAE using blurred images facilitates the generation of thin-line sketches (**Figure 4a**). We investigated the series of the generated images when the bottleneck state *z* was continuously changing (**Figure 4h**). The images generated by the *DS*_2_-trained VAE have two advantages compared with those trained by *DS*_1_: (1) when the change of *z* is not large so that the generated images look like the same digit, the images generated by *DS*_2_-trained VAE undergo more smooth and flexible shape morphing, whereas the images generated by *DS*_1_-trained VAE are more rigid; (2) when the change of *z* is large so that the generated images experience digit transition, the images generated by *DS*_2_-trained VAE look more realistic during the transition period, whereas the images generated by *DS*_1_ are more unrecognizable during the transition period (see Figure S2 for more examples). The smooth morphing of the generated images with *z* in *DS*_2_-trained VAE (**Figure 4h**) suggests that *DS*_2_-trained VAE develops a representation that respects the geometry of the images: similar images are represented by similar *z* values. Such geometry-respecting representation has been found in high-level brain areas such as prefrontal cortex and hippocampus, and may be important for cognitive flexibility of the animal [21, 22, 23]. Our results suggest that low-resolution top-down signals facilitates high-level brain areas to form such geometry-respecting representations.

In the computational model of V1 (**Figure 2**), we changed the coding resolution of the output of VAE decoder by adjusting *σ*_blur_ while keeping unchanged the input to VAE encoder to be the activities of V1 simple cells (**Figure 2a, right column**). In **Figure 4**, however, we used the same dataset (*DS*_1_ or *DS*_2_) as the input of VAE encoder and target output of VAE decoder during training, which means that the coding resolutions of the input and output of VAE are simultaneously changed. This design is because it is simpler to make the input and output of VAE be the same from the engineering aspect. In Figure S4, we studied the case when the input was kept to be *DS*_1_ while the target of the output was set to be *DS*_1_ or *DS*_2_. We found that the quality of the generated images is comparable with the case when the input is set to be the same as the target of the output. Therefore, it is the target of the output, instead of the input, that determines the quality of image generation.

Now we try to give explanations for the shape-morphing and input-irrelevance phenomena above from the perspective of complexity minimization, a fundamental principle in VAE. Complexity minimization assumes that the distribution *p*(*z|I*) = *N* (*µ, σ*) of bottleneck state *z* given a digital image *I* is close to the standard normal distribution *N* (0, 1) (**Figure 1a**). Shape morphing in **Figure 4h** can be understood from the following two soft constraints imposed by complexity minimization:

(1) First, any bottleneck state *z* near the center of *N* (0, 1) should generate a realistic-looking image through the VAE decoder: if a bottleneck state *z*_0_ close to the center of *N* (0, 1) cannot generate a realistic-looking image, the distribution *p*(*z|I*) of any image *I* should have a low value at the close-to-center point *z*_0_, violating the above assumption that *p*(*z|I*) is close to *N* (0, 1).
(2) Second, two similar bottleneck states correspond to similar decoder outputs. To understand this point, suppose *z*_middle_ is the middle point of *z*_1_ and *z*_2_ ( *z*_1_, *z*_2_ and *z*_middle_ are respectively transformed to three images *I*_1_, *I*_2_ and *I*_middle_ through the VAE decoder), then *I*_middle_ should look more like *I*_1_ than *I*_2_: if not, *p*(*z|I*_1_) will have two peaks at *z*_1_ and *z*_2_, and have a low value at *z*_middle_, violating the assumption that *p*(*z|I*_1_) is close to the unimodal function *N* (0, 1). This point can also be understood in another way: when training VAE, we sample the bottleneck state from a unimodal Gaussian distribution *N* (*µ, σ*), letting the output of the decoder converge to the same image *x* with different bottleneck states being sampled (**Figure 1a**). Therefore, two nearby bottleneck states are more likely to be sampled to train to output the same image; so after training, two nearby bottleneck states give similar outputs through the VAE decoder.

The natural solution to accommodate the above two soft constraints is that *I*_middle_ is a realistic-looking image interpolating *I*_1_ and *I*_2_, which is the shape-morphing phenomenon observed in **Figure 4h**. However, to make this interpolation successful, the images themselves should be smoothly interpolable: this is the advantage of low-resolution images, which is the case for *DS*_2_ (**Figure 4h, lower row**). When the images are not smoothly interpolable, unrealistic interpolation may happen, which is the case for *DS*_1_ (**Figure 4h, upper low**). To understand the input-irrelevance phenomenon (Figure S4), note that our argument above has nothing to do with the input: the job of VAE encoder is to transform the information in the input into the bottleneck state, which can be done as long as the input contains the information, no matter the representation of the input information is of low or high resolution. We will further illustrate the shape-morphing and input-irrelevance phenomena later using simplified models of visual processing, showing the importance of large receptive fields and low-spatial-frequency encoding of top-down signals (**Figure 6**).

To test the generalizability of the blurring technique (**Figure 4a**) above, we further examine the following situations:

(1) The generation of thin-line textures synthesized from OpenSimplex2 noise [31] (**Figure 4i**) using VAE.
(2) The generation of thin-line contours of the human faces in CelebA dataset [32] using VAE. We studied both constant-valued contours in which all pixels on the thin lines have the same intensity (see *DS*_1_ in **Figures 4b, i and 5a** for constant-valued contours) and continuous-valued contours in which different pixels on the thin line have different intensities indicating the intensity gradients of the corresponding CelebA image (Figure S6).
(3) The generation of constant-valued CelebA contours using generative adversarial networks (GAN), another dominating image-generation paradigm [33].

**Figure 5:**
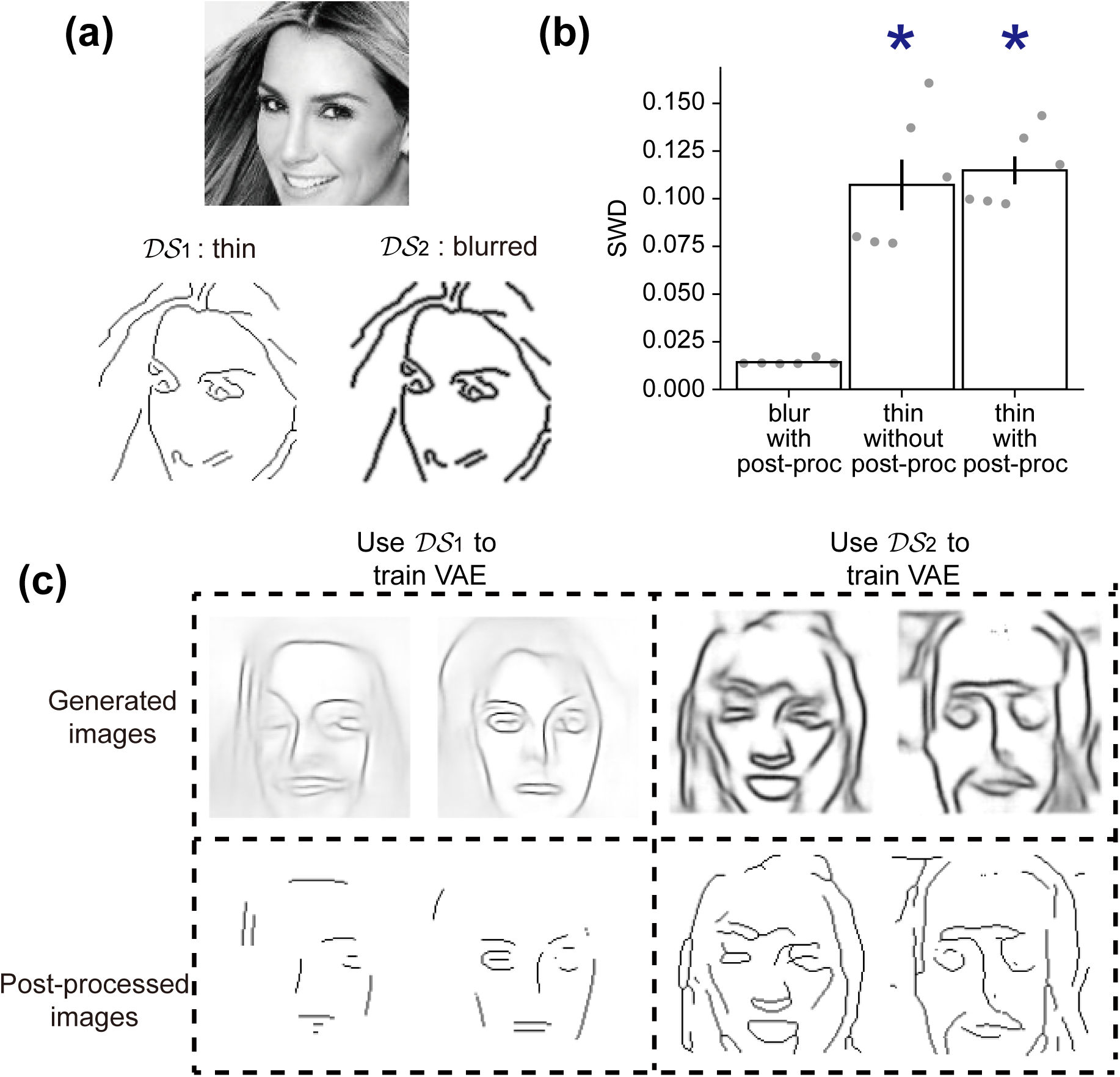
Low-resolution generation facilitates the generation of CelebA contours. (**a**) A gray-scale CelebA image (top), its thin contours (*DS*_1_, bottom left) and its blurred contours (*DS*_2_, bottom right). (**b**) Sliced Wasserstein distance (SWD) between the resulted images and the images in *DS*_1_ under different situations. Left bar: We train the VAE using blurred contours and post-process the VAE-generated images to infer the thin lines. Middle bar: We train the VAE using thin contours and do not post-process the VAE-generated images. Right bar: We train the VAE using thin contours and post-process the VAE-generated images. Error bars represent s.e.m. over 6 VAE configurations. Asterisks indicate *p <* 0.05 (Wilcoxon rank-sum test). (**c**) Examples of the VAE-generated (upper row) and post-processed (lower row) images, using *DS*_1_ (left column) or *DS*_2_ (right column) to train the VAE.

In all situations, the best way to generate thin-line sketches is to train VAE or GAN using blurred sketches, and then infer the thin lines from the generated images (**Figure 4a**), rather than train VAE or GAN using thin-line sketches directly (**Figure 5** and **Figures S5-S7**). These results confirm the generalizability of the proposed technique (**Figure 4a**).

### Low-resolution representations can be better simulated or reconstructed through top-down pathway

Overall, low-resolution top-down signals improves visual imagination and perception of the brain, and inspires a new AI technique to improve the generation of sketches. To get insight into the computational advantage of low-resolution top-down signals, we consider three toy models of V1 activity, where V1 neurons are located on a 1-dimensional line. In Model 1, only a single neuron has non-zero activity, while all the other neurons are silent (**Figure 6a, left**). In Model 2, the profile of neuronal activities is a Gaussian bump (**Figure 6a, middle**). In Model 3, this profile is an oscillating wavelet (**Figure 6a, right**). The shapes of the three profiles are parameterized by their positions on a one-dimensional line. Model 1 and Model 2 are respectively the correspondences of *DS*_1_ and *DS*_2_ (**Figure 4b**) in the one-dimensional case. We also consider Model 3 to better explain the mechanism.

**Figure 6:**
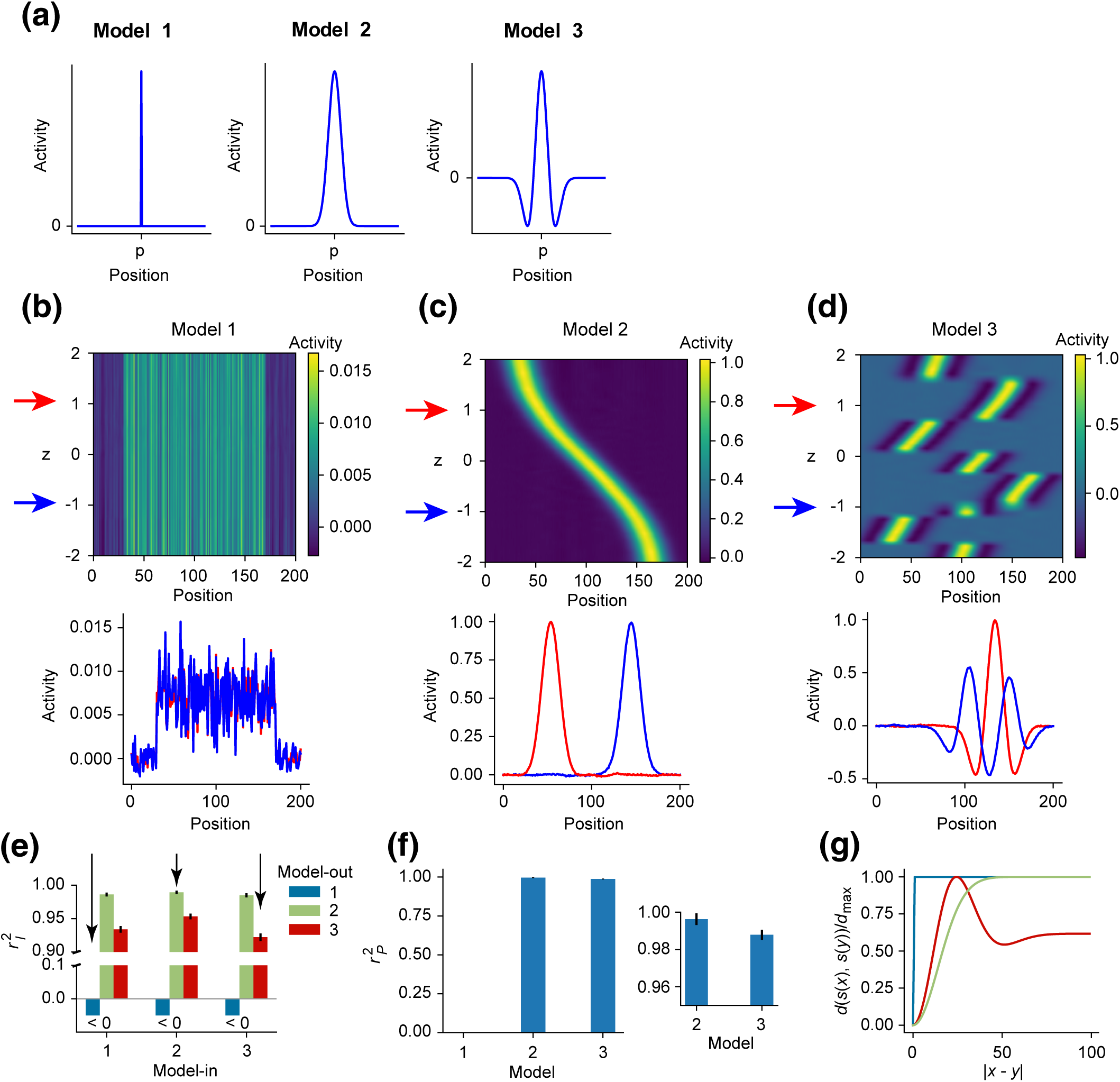
VAE is better at generating low-resolution representations. (**a**) Profiles of the three models of V1 activity. The peak (Model 1), bump (Model 2) and wavelet (Model 3) are parameterized by their positions *p* on a 1-dimensional line. (**b**) Upper panel: generated activity patterns as a function of the state *z* of the bottleneck variable. Lower panel: blue and red curves respectively represent the two generated patterns when *z* takes the two values (-1 and 1) indicated by the red and blue arrows in the upper panel. VAE is trained using the activity pattern of Model 1. (**c, d**) Similar to panel **b**, except that VAE is trained using the activity pattern of Model 2 (panel **c**) or 3 (panel **d**). (**e**) We let the input of VAE (i.e., *x* in Figure 1a) be the activity pattern of Model-in, but train the output of VAE (i.e., *x̂* in Figure 1a) to approach the activity pattern of Model-out. Both Model-in and Model-out can be Model 1, 2 or 3. We do not accurately show the value of 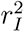 when 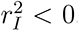. Arrows indicate the cases when Model-in equals Model-out, which are the cases in panels **b-d**. Error bars represent standard error of the mean (s.e.m.) over 1200 samples. (**f**) 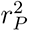 under the three models. Inset: results for Models 2 and 3 only. (**g**) Euclidean distance *d*(*s*(*x*)*, s*(*y*)) between two activity patterns *s* as a function of the distance *|x − y|* between the positions *x* and *y* of two dot stimuli on the 1-dimensional line, normalized by the maximal value of *d*, for Models 1 (blue), 2 (green) and 3 (red).

These three models can be biologically interpreted in various ways as the activities of different cell types in V1 in response to a dot stimulus on a 1-dimensional line:

(1) We can regard the neurons in all the three models as simple cells with different receptive-field sizes and spatial- frequency preferences. The receptive fields of the neurons in Model 1 have smaller sizes than those in Model 2. The receptive fields in Model 3 have the same spatial scale of Gaussian decay as the receptive fields in Model 2, but have higher spatial frequencies.
(2) The neurons in Model 2 can be regarded as the phase-insensitive complex cells downstream of the phase-sensitive simple cells in Models 1 and 3.

Therefore, by comparing Model 2 with Models 1 and 3, we can understand the computational advantage of large receptive-field size, low spatial-frequency tuning, and low grating-phase sensitivity in top-down signals in imagination and perception tasks [8, 9, 13].

We used each of the three models to train VAE, with the bottleneck state *z* of the VAE (**Figure 1a**) being one- dimensional. After training, we changed *z* in a range and investigated the output of the VAE decoder accordingly. In Model 1, sharp peaks are never generated by VAE, suggesting the failure of pattern generation (**Figure 6b**). In Model 2, Gaussian bumps are generated, with the bump position smoothly changing with *z* (**Figure 6c**). In Model 3, abrupt changes of wavelet positions sometimes happen with the continuous change of *z* (**Figure 6d**). These results suggest that generating low-resolution representations in Model 2 (i.e., the representations resulting from large receptive fields, low spatial-frequency preference and phase invariance) facilitates the high level representations of smooth shapes that can be morphed to fit or predict visual input (which is here the translational movement of the activity profiles along the 1-dimensional line).

We evaluated imagination quality by an index 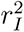, which indicates how well a VAE-generated pattern looks like an activity pattern of the model used to train the VAE. We evaluated perception quality by another index 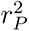, which indicates how well a VAE-generated pattern looks like the to-be-perceived pattern after iterative optimization (**Figure 1c**), see Methods . We found that both 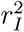 and 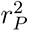 are maximal for Model 2, intermediate for Model 3, but is small (or even negative) for Model 1 (see the bars indicated by arrows in **Figure 6e**, also see **Figure 6f**). Therefore, the low-resolution representations in Model 2 can be better imagined or perceived. The disadvantage of Model 1 is apparent, because VAE trained by Model 1 failed to generate patterns like those in Model 1 (**Figure 6b**). The advantage of Model 2 over Model 3 is related to the smooth transition of *z* to the shape morphing (i.e., the translational movement) of the generated bumps (**Figure 6c**), because the generation quality is poor around the abrupt change points in Model 3 (see the blue curve in the lower panel of **Figure 6d** indicating the case when *z* = *−*1). The representation of stimuli by *z* in Model 2 is geometry-respecting, in the sense that two bumps spatially nearby are represented by similar *z* values, similar to **Figure 4h**.

In the study above, we input *P_a_* to VAE and trained VAE to generate *P_a_* in the output, with *P_a_* being the activity pattern of Model *a* (*a* = 1, 2, 3). Now we ask whether the advantage of Model 3 results from a ‘good’ pattern that the higher-level cortex receives from V1, or from a ‘good’ pattern that the higher-level cortex is to generate through the top-down pathway. To answer this question, we input *P_a_* to VAE but trained VAE to generate *P_b_* (*b ̸*= *a*). We found that the quality *r*^2^ of the generated images strongly depended on *P_b_*, but hardly on *P_a_* (**Figure 6e**). Therefore, the advantage of low-resolution representations is a top-down effect, and cannot be understood from a bottom-up perspective. We have mentioned a similar phenomenon before in the case of skeleton MNIST dataset (Figure S4).

To understand the reason for the advantage of Model 2, we studied the Euclidean distance *d*(*s*(*x*)*, s*(*y*)) between the activity patterns *s*(*x*) and *s*(*y*) as a function of *|x − y|*, where *s*(*x*) and *s*(*y*) are respectively the activity patterns stimulated by two dot stimuli on the 1-dimensional line at positions *x* and *y*. In Model 1, *d*(*s*(*x*)*, s*(*y*)) sharply jumps from 0 to a constant value at *|x − y|* = 0; in Model 3, *d*(*s*(*x*)*, s*(*y*)) is not monotonic; and in Model 2, *d*(*s*(*x*)*, s*(*y*)) monotonically and gradually increases with *|x − y|* (**Figure 6g**). This property of Model 2 is important for its advantage in top-down generation. To see this, suppose two nearby bottleneck states *z*_1_ and *z*_2_ (*z*_1_ *≈ z*_2_) generate two patterns *s*_1_ and *s*_2_, which correspond to two dot stimuli at positions *x*_1_ and *x*_2_ respectively. For illustration, we constrain that *s*_1_ is fixed during training, and *s*_2_ changes within the manifold *{s*(*x*)*}_x_*. VAE performs stochastic sampling to determine the bottleneck state, so the bottleneck state may be sampled to be *z*_2_ when *s*_1_ is provided as the input to the encoder and the target output of the decoder during training. By doing so, VAE tries to reduce the distance *d*(*s*_1_*, s*_2_) between the output patterns *s*_1_ and *s*_2_ that correspond to *z*_1_ and *z*_2_ respectively. In Model 2, due to the monotonicity of *d*(*s*_1_*, s*_2_) with *|x*_1_ *− x*_2_*|*, reducing *d*(*s*_1_*, s*_2_) reduces *|x*_1_ *− x*_2_*|*. In this way, two nearby bottleneck states *z*_1_ and *z*_2_ correspond to two nearby spatial positions *x*_1_ and *x*_2_ respectively; in other words, the bottleneck state represents the spatial geometry of the stimuli. In Model 3, *d*(*s*_1_*, s*_2_) is not monotonic with *|x*_1_ *− x*_2_*|*, so reducing *d*(*s*_1_*, s*_2_) does not always reduce *|x*_1_ *− x*_2_*|*, sometimes instead increases *|x*_1_ *− x*_2_*|*. In Model 1, *d*(*s*_1_*, s*_2_) remains constant when *s*_1_ = *s*_2_, so VAE has no gradient clue about how to update *s*_2_ close to *s*_1_.

This subsection shows that Gaussian bump signals (Model 2) can be better imagined or perceived than delta-peak signals (Model 1), which is consistent with our previous findings (**Figures 4 and 5**) in 2-dimensional cases. This subsection also shows that Gaussian bump signals (Model 2) outperforms oscillatory wavelet signal (Model 3). We also examined 2-dimensional cases and found similar results (**Figures S1-S5**).

### Sparse high-resolution information can be inferred from low-resolution representations

The previous subsection suggests that the higher-level cortex better simulates or reconstructs low-resolution representations for imagination or perception. For high-resolution information to be imagined or perceived, high-resolution information must be contained in such low-resolution representations. In other words, high-resolution information must be inferable from the top-down generated low-resolution representations. Here we study this coarse-to-fine inference problem.

We consider a population of low-coding-resolution neurons (**Figure 7a**) receiving stimuli from a one-dimensional circle. The feedforward connection from the stimulus receptor at position *θ_stim_* to the low-coding-resolution neuron preferring *θ_low_* Gaussianly decays with *|θ_stim_ − θ_low_|* (see Methods) . We study the following problem: given an activity pattern *P_low_* of the low-coding-resolution neurons, how to infer the corresponding pattern of the stimulus receptors?

**Figure 7:**
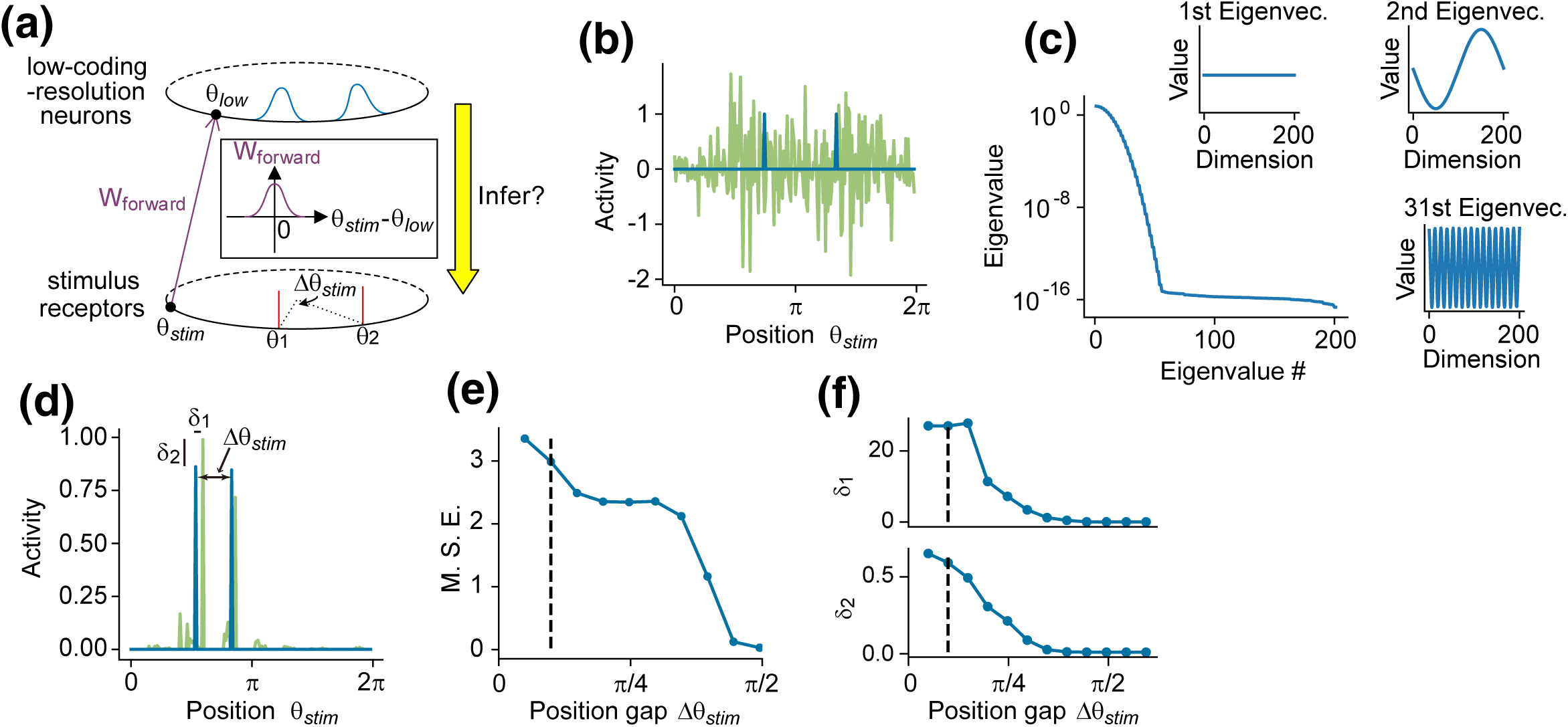
Sparse high-resolution information can be inferred from the activities of low-coding-resolution neurons. (a) Schematic of the model. Stimulus receptors and low-coding-resolution neurons are located on a circle. The feedforward connections ***W*** *_forward_* (magenta) from stimulus receptors to low-coding-resolution neurons Gaussianly decays with the difference *|θ_stim_ − θ_low_|* between their position preferences. The activities of stimulus receptors are delta-peaked (red) at *θ*_1_ and *θ*_2_ (with Δ*θ_stim_* = *|θ*_1_ *− θ*_2_*|*). The activities of low-coding-resolution neurons are Gaussianly bumped (blue). We study the inference of the activities of the stimulus receptors from the activities of the low-coding-resolution neurons (yellow arrow). (b) Blue: the activity of stimulus receptor as a function of *θ_stim_*. Green: inferred activity using 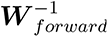 (**c**) Eigenspectrum of ***W*** *_forward_*. Insets are examples of eigenvectors. (**d**) The original activities (blue) of the stimulus receptors and the inferred activities (green) by a matching-pursuit algorithm. *δ*_1_ and *δ*_2_ respectively represent the differences in the positions and heights of the original and inferred peaks. (**e**) Mean-square error between the original and inferred activities as a function of the gap Δ*θ_stim_* between the positions of the two peaks. Vertical dashed line is the standard deviation of the Gaussian decay in ***W*** *_forward_*. (**f**) *δ*_1_ and *δ*_2_ as functions of Δ*θ_stim_*. In panels **e, f**, error bars representing s.e.m. over 8192 samples are smaller than the plot markers.

The most straightforward method lets the inferred stimulus be 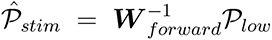, where ***W*** *_forward_* is the feedforward connection matrix from the stimulus receptors to the low-coding-resolution neurons. However, this method is unsuccessful (**Figure 7b**): ***W*** *_forward_* has very small eigenvalues corresponding to the eigenvectors that encode fine spatial details (**Figure 7c**), so 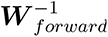 has very large eigenvalues. These large eigenvalues make 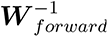 strongly amplify random noises (which were induced by limited numeric precision in **Figure 7b**), impairing the inference.

The inference difficulty results from the fact that a group *G* of stimulus receptors with similar position preferences give output to overlapping low-coding-resolution neurons, so that it is hard to tell which stimulus receptor causes an activity pattern of the low-coding-resolution neurons. If the stimulus receptors in *G* have sparse activities, so that in the ideal case only a single receptor in *G* is active, all the others being silent, then the inference can be achieved by exploiting the sparseness of *G*, similar to the compressed sensing scenario [55].

To evince the advantage of the sparseness of *G* for inference, we supposed that all stimulus receptors except those preferring two positions *θ*_1_ and *θ*_2_ were silent, and studied how the inference quality depended on Δ*θ_stim_* = *|θ*_1_ *− θ*_2_*|* (**Figure 7a**). When Δ*θ_stim_* is small, these two stimulus receptors have similar position preferences, so they can be considered to belong to the same group *G*; when Δ*θ_stim_* gets larger, these two stimulus receptors have farther different preferences, so the group *G* which contain stimulus receptors with similar position preferences have sparser activity.

We used a classical matching-pursuit algorithm [56] to perform this inference, which aims to explain a given pattern of low-coding-resolution neurons using as few active stimulus receptors as possible. We used three indexes to quantify the inference quality: (1) mean-square error of the inferred stimulus-receptor activity pattern *P*^^^*_stim_* from the original pattern *P_stim_*, (2) the difference *δ*_1_ between the positions of the peaks in *P*^^^*_stim_* and *P_stim_*, and (3) the difference *δ*_2_ between the heights of the peaks in *P*^^^*_stim_* and *P_stim_* (**Figure 7d**). All the three indexes indicated that the inference error was small when Δ*θ_stim_* was large (**Figure 7e, f**). These results suggest that sparse high-resolution information can be contained in low-resolution signals. In particular, those high-resolution features that cause similar activity patterns of low-coding-resolution neurons should have sparse strength.

Guided by the understanding of sparseness above, let us look back toward the V1 model (**Figure 2**) and the sketch- generation technique (**Figures 4, 5**). The sparseness in these investigations lies in the 1-pixel-width thin lines in the simple-cell activities (**Figure 2a, right column**) and the *DS*_1_ images (**Figures 4b, 5a**), which reduces the representation redundancy along the direction perpendicular to the thin line. Because of this sparseness, different lines can still be told apart after being blurred (see the *DS*_2_ image in **Figures 4b**), so that the original thin line can be inferred from the blurred lines: this is consistent with **Figure 7d-f**, where the inference of the two delta peaks is accurate when Δ*θ* is large enough so that the two Gaussian bumps are clearly separated apart.

## Discussion

In this paper, we show that top-down generation of low-resolution representations improves visual imagination and perception (**Figure 2**), and subsequently invent a simple but effective AI technique for sketch generation (**Figures 4 and 5**). The advantage of top-down low-resolution representations results from two reasons. First, low-resolution representations can be better simulated or reconstructed (**Figures 6**), due to its facilitation of the high-level cortex to form representations respecting to the geometry of stimuli (**Figures 6c and 4h**), consistent with the experimental observations in hippocampus and prefrontal cortex [21, 22, 23]. Second, sparse high-resolution information can be inferred from low-resolution representations (**Figure 7**). Therefore, top-down generation of low-resolution representations in V1 is not a shortcoming, but instead a strategy that the brain uses to improve visual imagination and perception.

Our analyses and numerical experiments all speak to different aspects of the analysis-by-synthesis and complexity- minimization principles inherent in the VAE model. This is manifested in terms of the sparse-coding principle of V1 [20] and has a close relationship with compressed sensing. From a machine-learning perspective, VAE is a neural-network implementation of a probability graph model (*z → x*), in which a latent variable *z* generates data *x*. In our sketch generation technique (**Figures 4, 5**), we implement a graph model (*z → x*_blur_ *→ x*_thin_) in which *x*_blur_ is blurred lines generated by VAE and *x*_thin_ is thin lines generated by the skeletonization operation (**Figure 4c**): similar coarse- to-fine technique has been explored in other machine-learning studies [54, 57, 58]. From a biological perspective (**Figure 2**), we propose that the brain implements a graph model (*z → x*_blur_), where *z* is the state of high-level brain areas, and *x*_blur_ is the blurry top-down representation to V1 (**Figure 2d**).

### The interaction between V1 and high-level brain areas

The information representation in V1 is sparse and high-dimensional [20]. In contrast, the representations in high- level areas such as prefrontal cortex and hippocampus are embedded in low-dimensional manifolds respecting the geometric structure of the knowledge [40, 41, 22]. Interestingly, when the brain performs working memory tasks, both V1 and high-level areas are active, with representations highly similar to when the brain perceives the remembered item [59, 6, 7]. Therefore, different representations of the same information exist in different brain areas. These representations can transform into each other, so the brain can leverage the advantages of different representations for better task performance. Our work suggests that top-down low-resolution signals may be a critical mechanism for establishing low-dimensional geometry-respecting representation in high-level brain areas (**Figures 6c and 4h**) and the transformation between the low-dimensional geometry-respecting representation in high-level brain areas and the sparse representation in V1.

Our work gains insight into the high-dimensional geometry of population responses in V1, with the *n*th principal component variance scaled as *n^−α^* with *α →* 1^+^ [60]. This geometry realizes the balance between efficient coding, which requires V1 neurons to code as much information using as few spikes as possible, and distance-respecting coding, which requires the distance between the V1 responses to two stimuli to increase with the distance between the two stimuli [60]. Efficient coding has been explained in feedforward models of V1 [61, 62]; here, we show that low-resolution feedback signals facilitate distance-respecting code (**Figures 6c, g and 4h**). Therefore, the *α →* 1^+^ scaling may result from the interaction between the bottom-up and top-down signals in V1. Experimentally, one may examine this hypothesis by studying whether *α* increases (i.e., V1 code gets whiter) when the animal becomes anesthetic so that top-down signals are disrupted. One may also study whether *α* decreases when the animal keeps the image in working memory with the eyes closed to disrupt the bottom-up signals.

From the perspective of the predictive-coding theory of perception [1], our model reveals that the activities of high-level areas are updated according to the error between the top-down signals and the low-resolution bottom- up representations of, possibly, complex cells, instead of the high-resolution representations of simple cells (**Figures 1c, d** and **2h**). In other words, the error signal is of low resolution. If this statement is correct, we should expect that the coding resolution of V1 neurons gets lower when the error signal is stronger: this is indeed the case when the visual stimulus is changing so that the mismatch between the real stimulus and the expectation of the brain is large. For example, the sizes of receptive fields expand when stimuli are moving [63], and the orientation and spatial frequency tuning of receptive fields is broad immediately after the onset of a stimulus [64, 65]. Therefore, the coding resolution of V1 neurons is low when the error signal is strong, which supports the low coding resolution of the error signal suggested by our work.

Our work leads to new interpretations of the computational role of complex cells. It is believed that the phase invariance of complex cells, which may stem from the convergent inputs from simple cells [66, 19], is to benefit object recognition [67, 68]. Some authors even proposed using alternate stacking of simple-cell and complex-cell layers to model the whole visual pathway [68, 69]. According to our work, however, the computational role of phase-insensitive complex cells is to provide simulation or reconstruction targets in imagination or predictive-coding perception (**Figure 1d**), facilitating the formation of geometry-respecting low-dimensional representations in high-level brain areas (**Figures 6c and 4h**).

One may argue that the low resolution of top-down signals to V1 is due to larger receptive fields of neurons in higher-level cortex (such as V2), resulting in lower resolution signals sent back to earlier layers. However, this explanation is not persuasive as the brain has plenty of mechanisms (such as feedforward or lateral inhibition [70]) to sharpen representations. Therefore, if low resolution led to inferior computational performance compared to high-resolution alternatives, top-down resolution should have increased over millions of years of evolution. Our study demonstrates the benefits of low-resolution top-down generation and suggests that evolution has favored this strategy for a reason.

### Inspiration to AI techniques

Sketches are ideal interfaces between human and computer software: we can easily modify sketches by drawing several lines, letting computers handle more tedious artistic work (such as image retrieval [71] or full-color image generation [72, 73]) based on the sketches we provide (see [74] for review). Sketch generation, the AI problem we study in this work, may further simplify the artistic interaction between humans and computers. However, compared with ordinary images, sketches are more challenging to generate, possibly due to their sparseness. There are mainly two approaches to generating sketches using deep networks. First is incremental generation. Instead of generating the whole image in a single step, we can generate sketches iteratively, one part after another. If sketches are represented by the coordinates of strokes in a vectorized format [75, 76, 77], sketches can be generated one stroke after another. If sketches are represented in a rasterized format [28, 78], sketches can be generated one image block after another. Second is deep-network aided loss-function definition. When training neural network models, instead of defining the loss function by directly using the difference between the generated and target sketches, we may input both the generated and target sketches into a deep network *N*, and define the loss function using the difference between the activities of *N* in response to the two inputs [79, 80, 81]. The blurring method introduced in this paper has its unique contribution compared to both approaches above. First, our method can improve the quality of the image generated by a single step, thereby reducing the number of iteration steps (thereby also the computational cost) to generate a high-quality image. Second, our method provides a simple yet effective definition of the loss function. By using blurred sketches, our method requires less computational cost than using a deep network to define the loss function. Our method is adaptable to both VAE and GAN (as shown in **Figures 4, 5** and Figure S7), indicating its versatility and broad applicability.

Sparse code with distance respect is a widely used AI technique that can improve the classification or generation of images [82, 83, 84]. Presently, the dominating approach for distance-respecting sparse code is based on local anchoring [85], representing image ***I*** as the linear combination of a subset of dictionary bases that have small Euclidean distances with ***I***. In our work, a thin line (which is sparse along the direction perpendicular to the thin line) is transformed into distance-respecting code through blurring (**Figure 6f**). This blurring idea may inspire future approaches for distance-respecting sparse code, which is an exciting research direction.

### Further discussions on the VAE model of the visual system

In this subsection, we will address three questions regarding our use of the VAE decoder to model the visual system. First, why do we use VAE (instead of other generative models such as GAN [33] and diffusion model [86]) to model the visual system? Second, compared with other common techniques for generating high-resolution images, what is the advantage of our method? Third, our neural network model is trained by gradient back-propagation (see Methods); however, back-propagation in the brain remains speculative [87], so whether our model is still valid?

(1) Here we use VAE decoder to model the top-down pathway from high-level brain areas to V1. There are other machine-learning generative models such as GAN [33] and diffusion model [86]. There are two reasons why we choose VAE. First is simplicity: GAN is more difficult to train than VAE due to instability problems [29], and diffusion model requires iterative noising/denoising operations which are computationally more expensive than VAE [86]. Second, perception is a two-stage process [5]: in the first stage, information is transmitted feedforwardly so that high-level brain areas can quickly form a rough estimation of the stimulus; in the second stage, this estimation is refined in a predictive-coding manner (**Figure 1c**). VAE is suitable to model the two-stage process of perception: given an image, the *µ* channel (see **Figure 1a**) is first calculated through the feedforward network of VAE encoder, then the initial state of the bottleneck state of the VAE decoder is set to be the value of *µ* channel before iterative predictive-coding updating. This is what we did in our numeric experiments (**Figures 3b, d, and 6e, f**) about perception (see Methods). On the contrary, GAN cannot model this first-stage feedforward perception process, and the encoder of diffusion model is a slow iterative noise-adding process, which is incompatible with the transient nature of the first-stage perception.
(2) Generating low-resolution images (i.e., not good at generating high-resolution details) is a usual tendency of VAE [29]. A common approach to improve the generation quality of VAE is to use hierarchical generative models, so that the resolution of the generative images is gradually improved at each hierarchical level [88, 58] (similar to the coarse-to-fine process of biological perception, in which the holistic gist of an image is perceived before fine local details [89, 90]), or use autoregressive priors in which the priors of VAEs are generated in a multi-stage manner [91]. Adding hierarchy is also a good practice to improve the generation quality of GAN [54], and is also the essential idea of diffusion models [86]. Compared with these hierarchical methods, the advantage of our method is lower computational cost. With the help of blurry representations, we can generate thin sketches using fewer neurons and less hierarchical computation, and the brain can also generate the information contained in V1 simple cells using fewer neurons and less hierarchical computation.
(3) Another potential issue is the network training method. We used gradient back-propagation (the most popular machine-learning method) to train our VAE model (see Methods). Though gradient back-propagation in the brain remains speculative [87], we do not think the training method undermines the validity of our conclusion (i.e., the computational advantage of low-resolution top-down signals). Numerous studies have shown the comparable dynamics between brain activities and the activities of artificial neural networks trained by gradient back-propagation (e.g., [92, 93]). Our recent study suggests that another machine-learning paradigm, evolutionary algorithm, can also train neural networks to exhibit similar dynamics to the brain [94]. Therefore, the dynamics may be irrelevant to the training methods, no matter whether the network is trained by gradient back-propagation, evolutionary algorithm in machine learning, or the biological plasticity rule in the brain. A possible explanation for this phenomenon is that different training methods universally drive the synaptic-weight configuration into a ‘high-entropy’ region in the weight-configuration space, so that if the weight configuration is slightly perturbed, the network still has good task performance [95, 96]: the similarity between high-entropy weight configurations may be the reason for the similarity of network dynamics resulting from different training methods.

## Methods

### The computational model of V1

#### Dataset preprocessing

We first converted CelebA images into grayscale, then cropped the 128 *×* 128 pixels around the center of the images, and then rescaled the images into 32 *×* 32 pixels.

#### The activities of V1 simple cells

To model the activities of V1 simple cells, we first got feature maps by filtering the preprocessed images using Gabor filters, then treated the feature maps by lateral WTA (**Figure 2a**).

There were 32 Gabor filters in total, with four orientations (0, 45, 90 and 135 degrees), two phases, and four spatial frequencies (0.18, 0.09, 0.045, 0.0225 cycle/pixel). The sizes were 4 *×* 4, 8 *×* 8, 16 *×* 16 and 32 *×* 32 for Gabor filters with spatial frequency 0.18, 0.09, 0.045, 0.0225 cycle/pixel respectively. For a preprocessed image *I*, we got two feature maps by taking the positive and negative parts corresponding to each Gabor filter *g*:

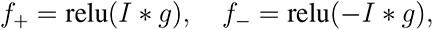

resulting in 64 feature maps in total. This Gabor filtering was performed using a customized python code modified from the ‘gabor’ routine of the scikit-image package.

For each feature map *f*, lateral WTA was performed by iterating the following sub-algorithm until the resulted feature map did not change anymore:

**Sub-algorithm:** For a pixel at position (*i, j*), we checked whether the intensity of this pixel was larger than all the pixels *P*_round_(*i, j*) within distance *d_W T A_* and 30*^◦^* from the direction of the Gabor filter corresponding to this feature map *f* (**Figure 2c**). If true, the intensities of *P*_round_(*i, j*) in *f* were set to zero. This operation above was iterated over all the positions (*i, j*).

The parameter *d_W T A_* above was 4, 8, 16, 32 pixels for Gabor filters with spatial frequency 0.18, 0.09, 0.045, 0.0225 cycle/pixel respectively.

The feature maps after lateral WTA modeled the activities of V1 simple cells and were the inputs of VAE.

#### The blurred representation

The training target of top-down signals to V1 (**Figure 2d**) was modeled by blurring the lateral-WTA-treated feature maps. The basic idea of this blurring was as follows. Suppose a pixel *p* had zero intensity in the lateral-WTA-treated feature map, we took another pixel *q* in the feature map. This *q* was the pixel spatially closest to pixel *p* with non-zero intensity. We then set the intensity of pixel *p* to be 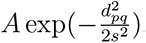. Here, *A* is the intensity of pixel *q*, *d_pq_* is the distance between *p* and *q*, and *s* = *σ*_blur_, 2*σ*_blur_, 4*σ*_blur_, 8*σ*_blur_ if the Gabor filter corresponding to the feature map has spatial frequency 0.18, 0.09, 0.045, 0.0225 cycle/pixel respectively. *σ*_blur_ is a free parameter to be adjusted in **Figure 3**.

#### The network structures of the VAE and post-processor

A variational auto-encoder (VAE) [18] contains an encoder and a decoder (**Figure 1a**). The encoder is a feedforward network that receives input ***x***_in_ and gives two channels of output ***µ*** and log(***σ***^2^). Then a vector ***z*** of random numbers is generated according to the Gaussian distribution *N* (***µ***, ***σ***), and is then inputted to the decoder. The decoder outputs ***x***^, which is required to approach ***x***_target_ during training. In the canonical definition of VAE (**Figure 1a**), the output ***x***^ is required to reconstruct the input (i.e., ***x***_target_ = ***x***_in_). This definition can be relaxed to the requirement that ***x***_target_ is treated from ***x***_in_ such that ***x***_target_ contains information of ***x***_in_. In **Figure 2e**, ***x***_target_ is ***x***_blur_, which is blurred from ***x***_in_ (i.e., ***x***_simple_), as illustrated in **Figure 2a, d**.

VAE is trained to minimize the following cost function:

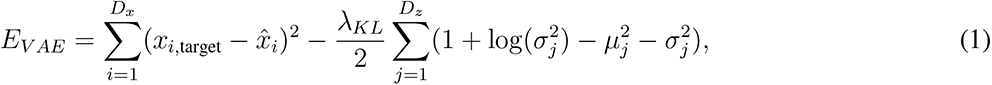

where *D_x_* is the dimension of the input and the output, *D_z_* is the dimension of the random variable ***z***. Minimizing the first term of this equation reduces the reconstruction error, minimizing the second term (which is the KL-divergence of *N* (***µ***, ***σ***) from the standard normal distribution *N* (**0**, **1**)) makes *N* (***µ***, ***σ***) close to *N* (**0**, **1**). *λ_KL_* is a parameter controlling the relative strengths of these two terms.

VAE was trained to generate the target top-down signals after receiving the activities of simple cells (**Figure 2e**), and then a post-processor was trained to infer the CelebA images from the output signals of VAE (**Figure 2f**). Both the VAE and post-processor were trained using the Adam optimizer [97] in Pytorch.

Both the VAE and post-processor were feedforward networks with input and output spatial size 32 *×* 32. The encoder of VAE had four hidden layers with spatial sizes 16 *×* 16, 8 *×* 8, 4 *×* 4 and 2 *×* 2 respectively. Similarly, the decoder of VAE also had four hidden layers with spatial sizes 2 *×* 2, 4 *×* 4, 8 *×* 8 and 16 *×* 16 respectively. The bottleneck state ***z*** of VAE had 32 dimensions. Adjacent layers in the encoder (decoder) were connected by convolutional (or transposed convolutional) kernels, followed by batch normalization and leaky ReLU activation function. The layers in the encoder had 64, 128, 256, 512 and 512 channels respectively, and the layers in the decoder had 512, 512, 256,128 and 64 channels respectively. *λ_KL_* = 0.5.

The post-processor was a U-Net [98], which is structurally similar to auto-encoder except that there is a direct link between the *i*th (counting from the input side) layer of the encoder and the *i*th (counting from the output side) layer of the decoder. The post-processor was to output realistic-looking images after inputting the output of VAE decoder. The encoder of the post-processor had an input layer with spatial size 32 *×* 32 and 64 channels, and also had four hidden layers with spatial sizes 16 *×* 16, 8 *×* 8, 4 *×* 4 and 2 *×* 2 and channel numbers 128, 256, 512, 512 respectively. The decoder of the post-processor had an output layer with spatial size 32 *×* 32 and 1 channel, and also had four hidden layers with spatial sizes 2 *×* 2, 4 *×* 4, 8 *×* 8 and 16 *×* 16 and channel numbers 512, 512, 256, 128 respectively. The bottleneck state of the U-Net had 32 dimensions. Adjacent layers in the encoder (decoder) were connected by convolutional (or transposed convolutional) kernels, followed by batch normalization and leaky ReLU activation function.

To train the post-processor, we inputted the simple cell activities ***x***_simple_ into the VAE encoder, and let the bottleneck state of the VAE be the *µ*-channel of the encoder, then passed the output images of VAE through the post-processor, getting ***x***_post-proc_ (**Figure 2f**). We adjusted the synaptic weights in the post-processor by minimizing the mean-square error (MSE) loss function plus an adversarial loss

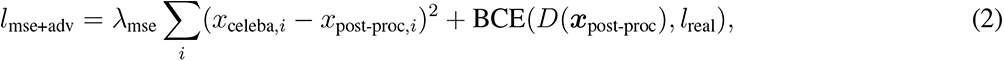

where *x*_celeba_*_,i_* and *x*_post-proc_*_,i_* are the *i*th pixel in ***x***_celeba_ (i.e., preprocessed CelebA image) and ***x***_post-proc_ respectively, *λ*_mse_ = 0.01, and BCE(*D*(***x***_post-proc_)*, l*_real_) is the binary cross-entropy between the output *D*(***x***_post-proc_) of an adversarial network in response to ***x***_post-proc_ as input and the label *l*_real_ = 1 for real images.

The adversarial network was also a feedforward network. Its input had spatial size 32 *×* 32 and 1 channel, ready to receive inputs from the output of the post-processor for discrimination. The output of the adversarial network had spatial size 1 *×* 1 and 1 channel. It had 3 hidden layers with spatial sizes 16 *×* 16, 8 *×* 8 and 4 *×* 4 with channel numbers 16, 32 and 64 respectively. Adjacent layers were connected by convolutional kernels, followed by batch normalization and leaky ReLU activation function. The output was treated by the sigmoid activation function.

The parameters in the adversarial network were updated by minimizing

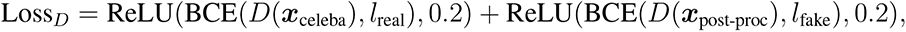

where ***x***_celeba_ indicates a CelabA image after preprocessing, *l*_real_ = 1 is the label for real images, *l*_fake_ = 0 is the label for fake images, ReLU(*x,* 0.2) equals *x* if *x >* 0.2 but equals zero otherwise. We introduced this ReLU(*x,* 0.2) function here to stop training the adversarial network if the adversarial network is already powerful enough, which improves the stability of GAN.

#### The quality of imagination and perception

Imagination was modeled by sampling the bottleneck state from standard normal distribution (**Figure 2g**), getting the output of the VAE decoder and post-processing the VAE output. The imagination quality (**Figure 3c**) was quantified using sliced Wasserstein distance (SWD) [53, 54] between the post-processed VAE-generated images ***x***_post-proc_, and the preprocessed CelebA images ***x***_celeba_. SWD is defined by

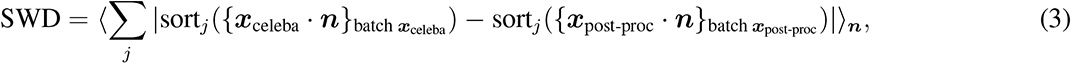

where ***n*** is a randomly selected direction vector with ∥***n***∥^2^ = 1, sort*_j_* (*{****x***_celeba_ *·* ***n****}*_batch_ ***_x_***_celeba_ ) indicates the *j*th element after sorting the value ***x***_celeba_ *·* ***n*** in a batch of ***x***_celeba_, ⟨·⟩***_n_*** means averaging over ***n***. Numerically, the batch sizes of both ***x***_celeba_ and ***x***_post-proc_ were 1024, and we averaged over 512 directions of ***n*** when calculating SWD.

Perception is a two-stage process: information is transmitted feedforwardly at the early stage and processed recurrently at the later stage, reflecting the shift of computational demand from responding speed to representation resolution [5]. We modeled the early stage by inputting the simple cell activities (**Figure 2a, right column**) into the VAE encoder, then setting the bottleneck state of the VAE to be the *µ*-channel of the encoder. We modeled the later stage as apredictive-coding process [1] (**Figure 2h**), in which the bottleneck state *z* was updated to minimize the error between the top-down representation ***x***_tp_ and the blurred representation ***x***_blur_ (**Figure 2d, bottom row**), after we initialized *z* to be the *µ*-channel of the encoder in the first stage before the iterative updating.

The perception quality (**Figure 3d**) was quantified in the following way. Given a preprocessed CelebA image ***x***_celeba_, we inputted the lateral-WTA-treated feature maps ***x***_simple_ (**Figure 2a, right column**) into the VAE encoder. We set the initial bottleneck state of the VAE decoder to be the *µ*-channel of the VAE encoder in response to the lateral- WTA-treated feature maps, and then optimized the bottleneck state to minimize the mean square error between the blurred feature maps (**Figure 2d**) and the output of the VAE decoder (see **Figure 2h**). The perceived image ***x***_post-proc_ was then inferred by passing the output of the VAE decoder through the post-processor. The perception quality was quantified using an index *U* (**Figures 2h, 3d**) defined as

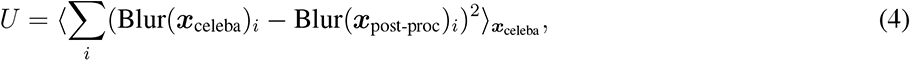

where Blur(*I*)*_i_* means the *i*th pixel of the image *I* after blurred by a Gaussian filter of 1-pixel width. This blurring operation was performed to reflect the contribution to *U* by fine-spatial details. To understand this, suppose there are two 1-pixel-width lines taking the same direction in ***x***_celeba_ and ***x***_post-proc_ respectively. In this case, Σ*_i_*(***x***_celeba_*_,i_ −****x***_post-proc_*_,i_*)^2^ is zero if these two parallel lines overlap, whereas takes the same positive value if these two parallel lines do not overlap, no matter whether they are located spatially nearby or far apart. By blurring ***x***_celeba_ and ***x***_post-proc_, *U* gradually increases with the distance between these two parallel lines, which is consistent with the intuition that the perception error increases with the distance between these two lines.

### The generation of skeleton MNIST images

#### VAE structure

In the VAE used in **Figure 4**, the encoder is a multilayer perceptron (MLP) which has an input layer of size 28 *×*28 = 784 and three hidden layers of sizes 512, 256 and 128 respectively. The bottleneck state has dimension 20. The decoder of the VAE is also a MLP which has three hidden layers of sizes 128, 256 and 512 respectively and an output layer of size 28 *×* 28 = 784. Adjacent layers are all-to-all connected. Leaky ReLU is the activation function. The loss function is defined in **eq. 1**.

#### The three datasets of 2-dimensional images

The dataset *DS*_1_ in **Figure 4b** is the skeleton MNIST dataset [30]. The intensity of a pixel in an image in *DS*_1_ is 1 or 0 depending on whether this pixel belongs to a line of 1-pixel width.

The dataset *DS*_2_ was blurred from *DS*_1_ using the following method. Given a pixel *p*_0_ not belonging to thin lines, we looked for a pixel *p*_1_. This *p*_1_ should belong to thin lines, and at the same time are spatially closest to *p*_0_. We then set the intensity of *p*_0_ to be exp(*−d*(*p*_0_*, p*_1_)^2^*/*2), where *d*(*p*_0_*, p*_1_) indicates the spatial distance between the pixels *p*_0_ and *p*_1_.

In **Figures S1-S4**, we also consider another dataset *DS*_3_, as the 2-dimensional counterpart of Model 3 in **Figure 6a**. *DS*_3_ was treated from *DS*_1_ in a similar way to *DS*_2_, except that the intensity of *p*_0_ was set to be *a* exp(*−d*(*p*_0_*, p*_1_)^2^*/*2), where *a* = *−*1 if *d*(*p*_0_*, p*_1_) = 1 and *a* = 1 otherwise.

#### The quality of image generation

The images generated by VAE were post-processed in two steps (**Figure 4c**). First, images were binarized such that pixels with intensities larger (or smaller) than a threshold *θ_thres_* were set to 1 (or 0). Second, the images were skeletonized using the ‘skeletonize’ routine of the skimage python package.

To quantify the quality of the post-processed images, we trained an MLP to classify the skeleton MNIST dataset (i.e., *DS*_1_). This MLP contained a hidden layer of size 1000 with a leaky-ReLU activation function. After receiving a post-processed VAE-generated image *I*, this MLP output a label distribution *p*(*x|I*) (*x* = 0, 1*, · · ·,* 9). In **Figure 4e**, *H_realistic_* = *E_I_* [− Σ*_x_ p*(*x|I*) ln *p*(*x|I*)], where *E_I_* [*·*] means average over all post-processed VAE-generated images [99]; in **Figure 4f**, *H_xcat_* = − Σ*_x_ E_I_* [*p*(*x|I*)] ln *E_I_* [*p*(*x|I*)] [99]. To plot **Figure 4g**, we first chose the post-processed VAE-generated images with high realisticity (i.e., max*_x_ p*(*x|I*) *>* 0.9), then for all the images belonging to a category *x*, we calculated the variance *λ_i_*(*x*) along the *i*th principal component (PC), *D_incat_* was defined as 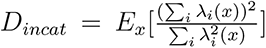. **Figure 4e-g** shows how *H_realistic_*, *H_xcat_*, *D_incat_* change with the binarization threshold *θ_thres_* and the parameter *λ_KL_* in **eq. 1**. Note that if *θ_thres_* is high, the image after post-processing may be very sparse (i.e., only a few pixels are nonzero), especially when *λ_KL_* also takes a large value. In this case, the MLP network has an artifact that *p*(*x|I*) strongly peaks at *x* = 1, and *p*(*x*≠ 1*|I*) has a very small value. Because of this artifact, in **Figure 4e-g**, we excluded the data points at which the percentage of nonzero pixels in the post-processed images was smaller than 1%. Some data points when *λ_KL_* = 0.9 for *DS*_1_ resulted in images with sparsity a little larger than 1%, but we also excluded these data points, because the quality of the generated images was really bad. This artifact was weak for *DS*_2_ in our range of parameters, so we plotted the whole parameter range for *DS*_2_. Because of the same reason, we also excluded the data for *DS*_3_ when *λ_KL_* = 0.5, 0.7, 0.9 in **Figures S1c-e, S3 and S4c-e**.

**Figure 4h** was plotted by gradually and linearly changing the bottleneck states of VAEs from ***z*** = [1.5, 1.5*, · · ·,* 1.5] to [*−*1.5*, −*1.5*, · · ·, −*1.5].

### The generation of CelebA contours

#### Preparing the contour dataset

We converted CelebA images into gray-scale, and cropped the resulted gray-scale images around the center, resulting in 128 *×* 128 image patches. Then, we normalized the mean of each patch to be the mean of all the obtained patches. To detect contours from the normalized gray-scale patches, we followed the standard technique (see https://learnopencv.com/edge-detection-using-opencv/): we first blurred the patches using a Gaussian filter with astandard deviation of 2 pixels, then used the ‘Canny’ routine of OpenCV (with parameters threshold1=50, threshold2=100, L2gradient=True) to obtain binary contours. We then skeletonized the binary contours using the ‘skeletonize’ routine of the skimage package, resulting in binary contours of 1-pixel width, which are images in dataset *DS*_1_ (**Figure 5a**). Images in dataset *DS*_2_ were obtained by blurring images in *DS*_1_, similar to the way to process skeleton MNIST images introduced above.

#### The network structure of VAE

The VAEs to generate CelebA contours were deep feedforward neural networks with input and output spatial size 128 *×* 128. The encoder of VAE had six hidden layers with spatial sizes 64 *×* 64, 32 *×* 32, 16 *×* 16, 8*×* 8, 4 *×* 4 and 2 *×* 2 respectively. Similarly, the decoder of VAE also had six hidden layers with spatial sizes 2 *×* 2, 4 *×* 4, 8 *×* 8, 16 *×* 16, 32 *×* 32 and 64 *×* 64 respectively. The bottleneck state of VAE had 128 dimensions. Adjacent layers in the encoder (decoder) were connected by convolutional (or transposed convolutional) kernels, followed by batch normalization and leaky ReLU activation function. The layers in the encoder had 1, 64, 128, 256, 512, 512 and 512 channels respectively, and the layers in the decoder had 512, 512, 512, 256, 128, 64 and 1 channels respectively. *λ_KL_* = 0.5.

#### Post-processing the VAE-generated images

VAE-generated images were post-processed similarly to that of **Figure 4c**, with binarization threshold *θ_thres_* = 0.35.

#### Evaluating the quality of contour generation

The quality of contour generation was quantified by the SWD (**eq. 3**) between the post-processed VAE-generated images and the images in *DS*_1_ (the left and right bars in **Figure 5b**). To plot the middle bar in **Figure 5b**, we also calculated the SWD between the images generated by *DS*_1_-trained VAEs without post-processing and the images in *DS*_1_.

### The imagination and perception of 1-dimensional images

#### VAE structure

In the VAE used in **Figure 6**, the encoder is a multilayer perceptron (MLP) which has an input layer of size 201 and three hidden layers of sizes 100, 50 and 20 respectively. The bottleneck state has dimension 1. The decoder of the VAE is also a MLP which has three hidden layers of sizes 20, 50 and 100 respectively and an output layer of size 201. Adjacent layers are all-to-all connected. Leaky ReLU is the activation function.

#### The three models of 1-dimensional images

In the three models in **Figure 6a**, 201 neurons are positioned on a one-dimensional line. In Model 1, the activity pattern is a delta peak *f*_1_(*x*; *a*) = *δ_x,a_* in which only a single neuron at position *a* has activity 1, whereas all the other neurons have zero activity. In Model 2, this pattern is a Gaussian bump 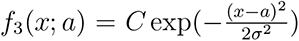, where *σ* = 10, *C* is a normalization factor such as max*_x_ f*_3_(*x*; *a*) = 1. In Model 3, this pattern is a Gabor wavelet *f*_2_(*x;a*) = 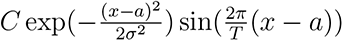, where *σ* = 10, *T* = 80, *C* is a normalization factor such that max*_x_ f* (*x*; *a*) = 1. In *f*_1_(*x*; *a*), *f*_2_(*x*; *a*) and *f*_3_(*x*; *a*), *a* is a random integer from 31 to 171.

#### The quality of imagination

To quantify the quality of the generated patterns (**Figure 6e**), for any generated pattern *p*, we defined 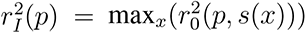, where *s*(*x*) is the activity pattern of Model-out in response to the dot stimulus at position *x* on the 1-dimensional line, and 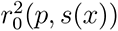 is the ratio of the variance of *p* that can be explained by *s*(*x*) (i.e., coefficient of determination). Specifically, 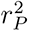(*p,s*) is defined by

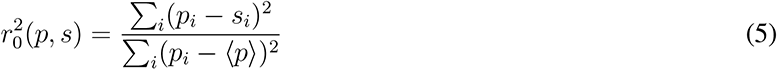

where *p_i_* and *s_i_* are respectively the activities of the *i*th neuron in the patterns *p* and *s*, *⟨p⟩* is the mean activity of pattern *p*.

#### The quality of perception

We examined the quality of perception using the index 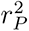 (**Figure 6f**), which was calculated as follows:

We set the target image *T* (i.e., blue curve in **Figure 6c**) to be an activity pattern of Model *a* (*a* = 1, 2, 3), and initialized the bottleneck state to be the value of the *µ*-channel (see **Figure 1a**) of the VAE encoder when inputting *T* into the encoder. The value of the *µ*-channel is the best estimation of the optimal bottleneck state by the encoder. Then we updated the bottleneck state to minimize the error between *T* and the decoder output using Adam optimizer [97]. We defined *r*^2^ as the coefficient of determination to quantify how much variance of the target image can be explained by the generated image.

### Inferring high-precison information from the activities of low-coding-resolution neurons

In our simulation for **Figure 7a**, both the stimulus receptor and the low-coding-resolution neuron layers contained

*N* = 201 neurons lying on a one-dimensional circle. Only two stimulus receptors were active, separated by

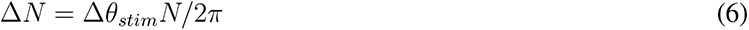

neurons, with Δ*N* = 5, 10, 15*, · · ·,* 60. The activities of the two active stimulus receptors were set to two random values in the range [0.5, 1], and the activities of all the other stimulus receptors were zero.

The feedforward connection from a stimulus receptor preferring *θ_stim_ ∈* [0, 2*π*) to a low-coding-resolution neuron preferring *θ_low_ ∈* [0, 2*π*) was

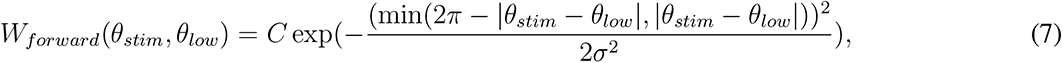

where *σ* = 2*π ×* 10*/N*, *C* is a normalization factor so that 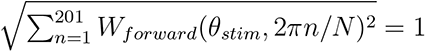.

Given the activities *{x*_1_*, x*_2_*, · · ·, x_N_ }* of stimulus receptors, we calculated the activities *{y*_1_*, y*_2_*, · · ·, y_N_ }* of low- coding-resolution neurons using the feedforward matrix ***W*** *_forward_*, and then we studied how to infer the activities *{x̂*_1_*, x̂*_2_*, · · ·, x̂_N_ }* of stimulus receptors using *{y*_1_*, y*_2_*, · · ·, y_N_ }*.

We tried a matching-pursuit algorithm [56] in **Figure 7d-f**. Specifically, we first defined the remnant *r_i_* = *y_i_*, and set *x̂_i_* = 0 (*i* = 1, 2*, · · ·, N* ), then iterated the following steps for 1000 times:

(i) Calculate 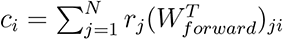, then set *c_k_* = 0 *k* = arg max_*i*_ |*c_i_*.
(ii) Let 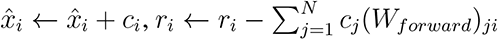

## Acknowledgments

We thank Prof. Yuanyuan Mi, Prof. Changsong Zhou and Prof. Shuzhi Sam Ge for comments on the manuscript and helpful discussions. Z.B. is supported by the National Natural Science Foundation of China (Grant No. 32000694).

## Supplementary materials

### 1 More results on skeleton MNIST images

See **Figures S1-S3** for more results on skeleton MNIST images.

### 2 Generating thin-line textures using VAE

We trained VAE using textures treated from OpenSimplex2 textures (**Figure S5a**) with various image sizes and *λ_KL_* values. We found that the thin-line textures inferred from the VAE-generated images have the best quality if the VAE was trained by *DS*_2_ (**Figure S5b**). This advantage is demonstrated by the small sliced Wasserstein distance (SWD) between the post-processed images generated by *DS*_2_-trained VAE and the *DS*_1_ dataset under a broad parameter range, especially when the image size is large (**Figure S5c-e**). These results again demonstrate the advantage of training VAE using blurred images to generate thin-line sketches.

#### Image preparation

The OpenSimplex2 textures were generated using the FastNoiseLite package with the following parameters. Noise type: OpenSimplex2; frequency: 0.05; fractal type: ridged; fractal octaves: 1; fractal lacunarity: -0.29; fractal gain: 0.05; fractal weighted strength: 1.3; domain warp type: OpenSimplex2; domain warp amplitude: 5. The generated images were then binaried through threshold 0.5, and skeletonized by the ‘skeletonize’ routine of skimage python package, resulting in 60000 images of size 64 *×* 64.

To plot **Figure S5c-e**, we extracted image patches of different sizes (8 *×* 8, 16 *×* 16, 32 *×* 32 and 64 *×* 64) at random positions. For each image patch, we set the intensities of the pixels whose distances with boundaries were 1 or 2 to be zero, resulting in dataset *DS*_1_. Datasets *DS*_2_ and *DS*_3_ were derived from *DS*_1_, similar to those for skeleton MNIST images.

#### The network structure of VAE

The structure of the VAE to generate textures was the same as the structure of the VAE we used to generate CelebA contours (**Figure 5**), see Methods . *λ_KL_* = 0.5.

**Figure S1:**
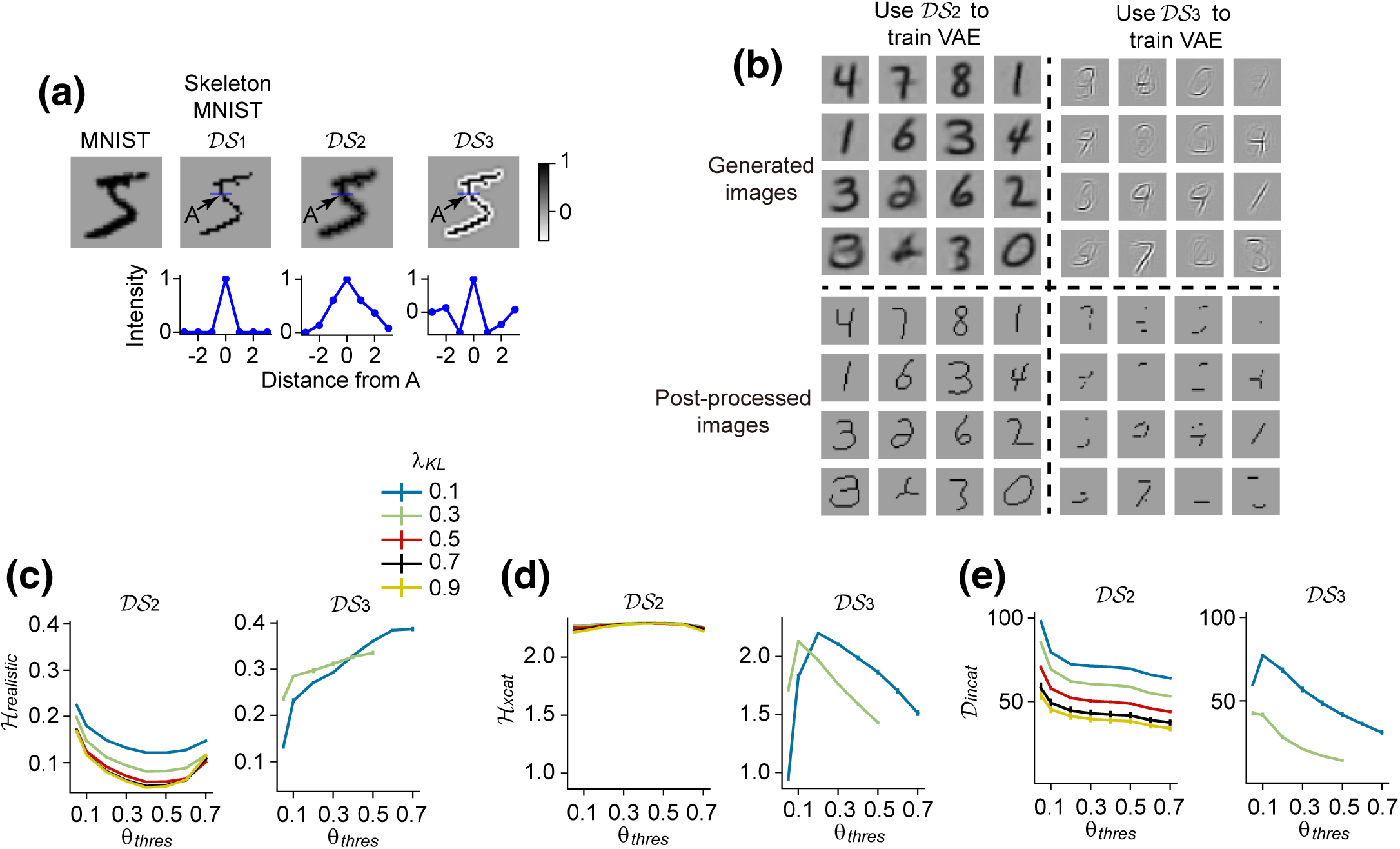
The ‘oscillatory’ dataset and the quality of VAE generation when VAE is trained by this dataset. (a) Illustration of the two datasets *DS*_1_ and *DS*_2_ that already appear in the main text and another new dataset *DS*_3_. Similar to Figure 4b in the main text. In *DS*_1_, *DS*_2_ or *DS*_3_, the pixel intensities delta peak, Gaussianly decay or oscillating decay along the direction perpendicular to a line respectively, so that *DS*_1_, *DS*_2_ or *DS*_3_ are regarded as the displacement of Model 1, 2 or 3 (see Figure 6a) respectively for the 2-dimensional-image case. (**b**) Upper subplots: examples of the generated images when using datasets *DS*_2_ and *DS*_3_ to train VAE. Lower subplots: the generated images after post-processing. Note that *DS*_2_ results in better image quality than *DS*_3_. (**c-e**) *H_realistic_*, *H_xcat_* and *D_incat_* as functions of the binarization threshold *θ_thres_* when the parameter *λ_KL_* in VAE (see Methods) takes different values, when VAE is trained using *DS*_2_ and *DS*_3_ respectively. Some data points for *DS*_3_ are missing, see Methods for explanation.

**Figure S2:**
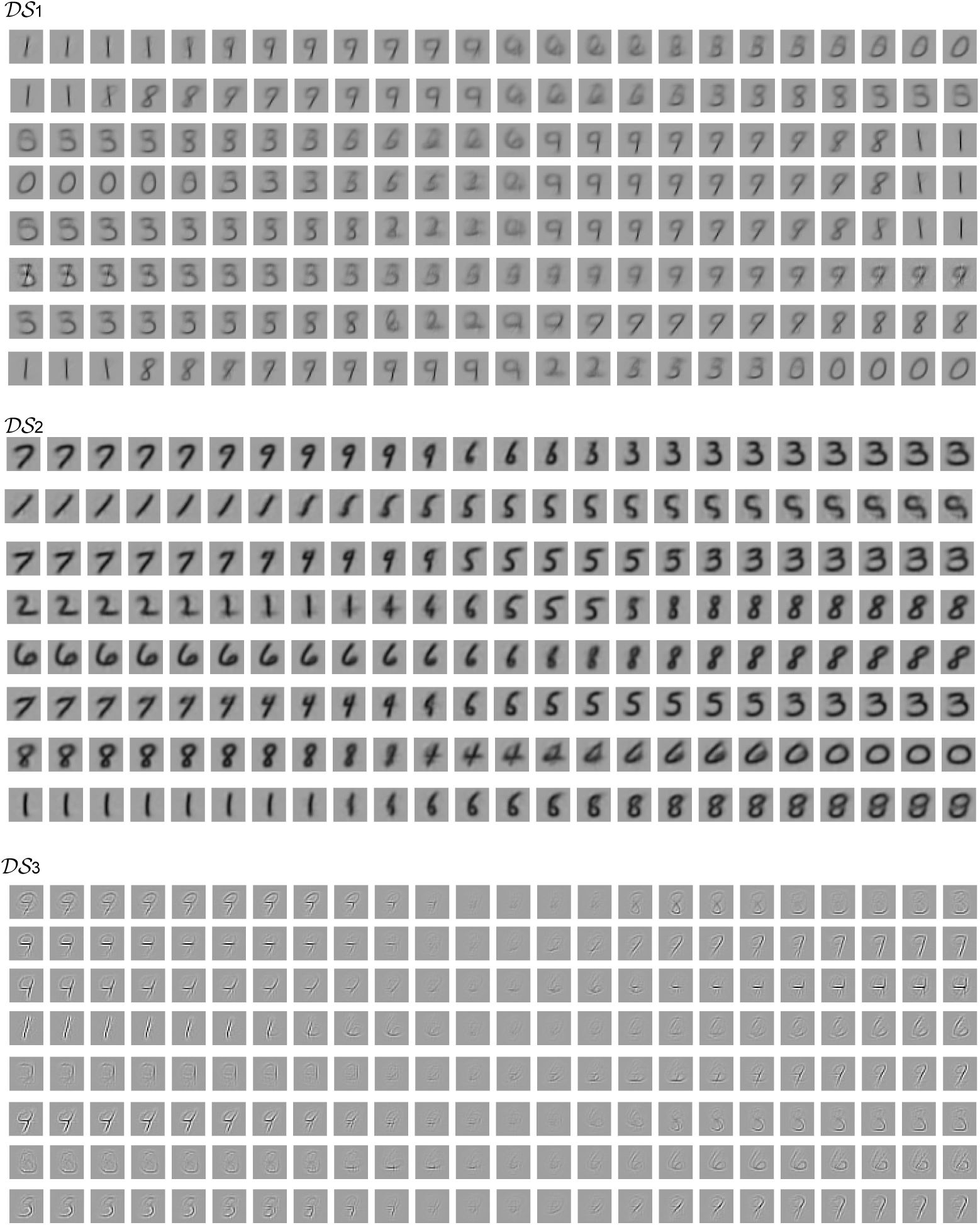
More examples of the generated images of the VAEs trained by the three datasets. *DS*_1_, *DS*_2_ **and** *DS*_3_ **respectively.** Each row represents the images generated by a VAE configuration when the bottleneck state slowly and linearly changes from ***z*** = [1.5, 1.5*, · · ·,* 1.5] to [*−*1.5*, −*1.5*, · · ·, −*1.5]. *λ_KL_* = 0.7 for both *DS*_1_ and *DS*_2_, *λ_KL_* = 0.3 for *DS*_3_.

**Figure S3:**
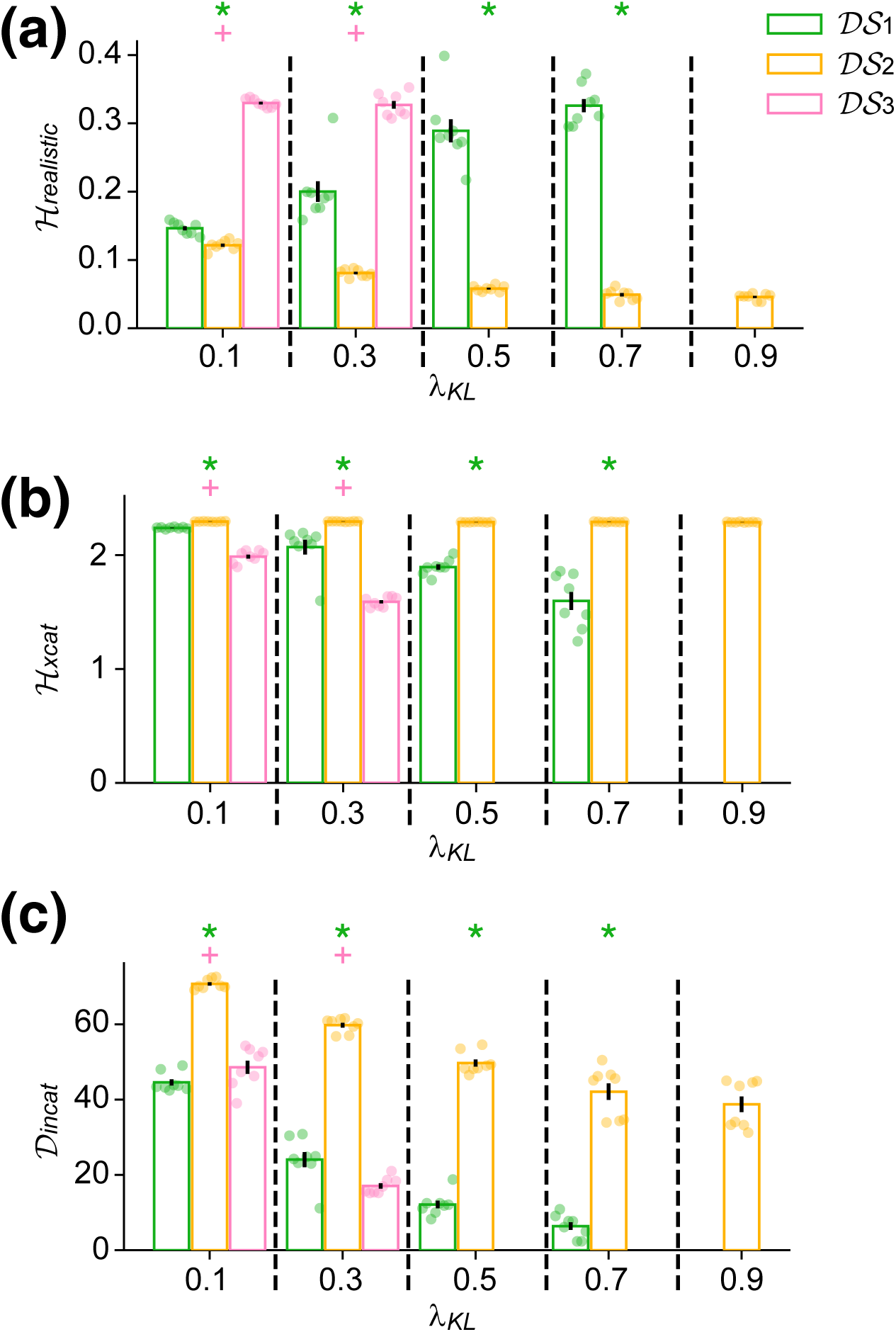
More data on the generation of skeleton MNIST images. Realisticity entropy *H_real_* (panel **a**), cross-category entropy *H_xcat_* (panel **b**) and within-category dimensionality *D_incat_* (panel **c**) as functions of *λ_KL_* when the binarizing threshold *θ_thres_* = 0.4. Each dot represents the result of a single VAE configuration. Error bars represent s.e.m. over 8 VAE configurations. Vertical dashed black lines separate data of different *λ_KL_* values. Some data are missing due to the same reason for the missing data in Figure 4e-g in the main text, see Methods in the main text. In panel **a**, green asterisk (or blue plus sign) means that *H_real_* for *DS*_2_ is significantly (*p <* 0.05) smaller than that for *DS*_1_ (or *DS*_3_) at the same *λ_KL_*. In panel **b**, green asterisk (or blue plus sign) means that *H_xcat_* for *DS*_2_ is significantly larger than that for *DS*_1_ (or *DS*_3_). In panel **c**, green asterisk (or blue plus sign) means that *D_incat_* for *DS*_2_ is significantly larger than that for *DS*_1_ (or *DS*_3_).

**Figure S4:**
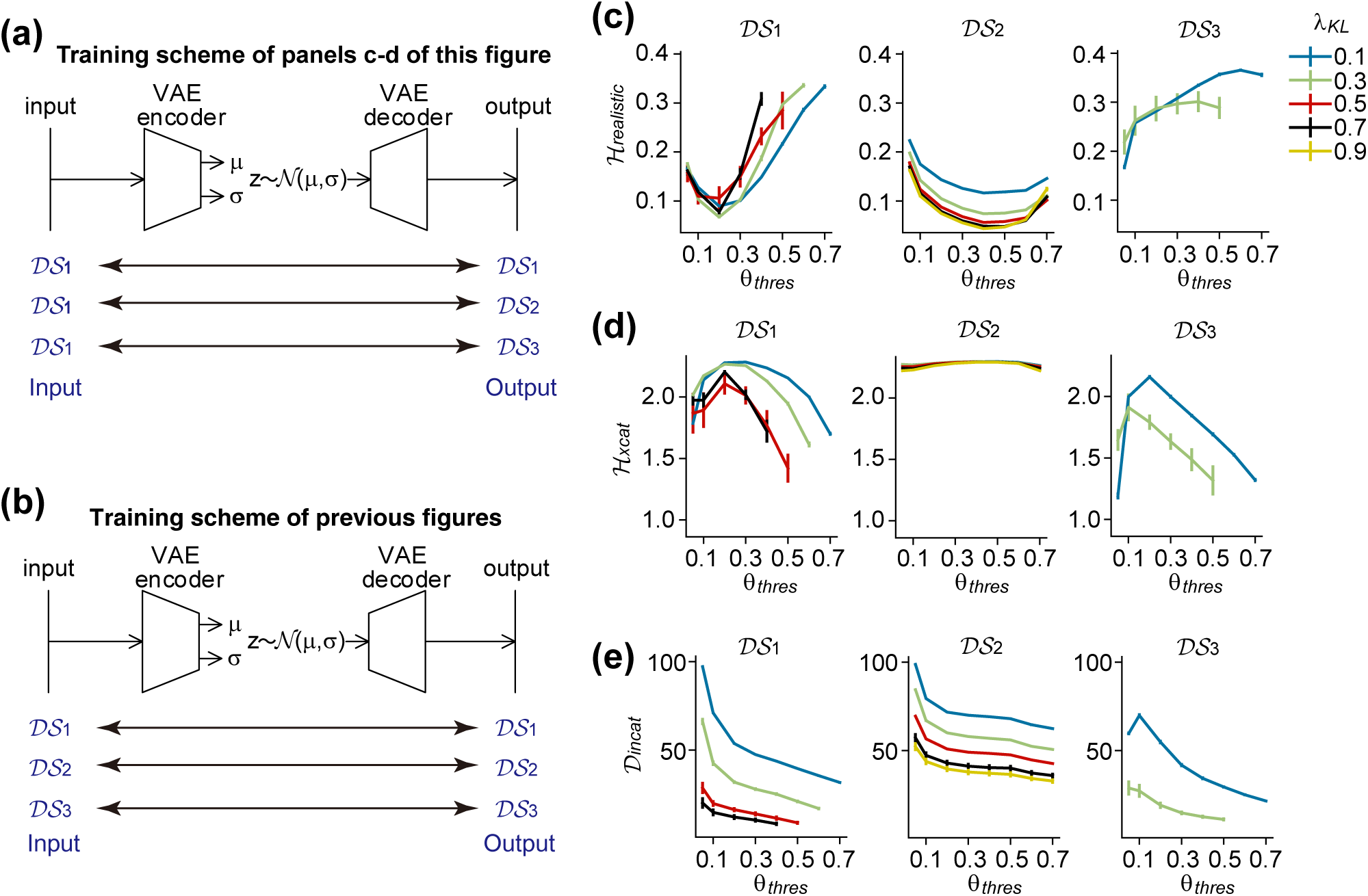
The quality of the VAE-generated images after post-processing when the VAE encoder always receives *DS*_1_ images during training. (a) The training scheme of panels c-d in this figure. We always inputted *DS*_1_ to the VAE encoder, but trained the decoder output to approach the corresponding images in *DS*_1_, *DS*_2_ or *DS*_3_. A double arrow indicates that ‘corresponding’ images in the two datasets at the two ends of the double arrow are simultaneously used to train a VAE. By ‘corresponding’, we mean that two images in two datasets *DS_x_* and *DS_y_* (*x, y* = 1, 2, 3) are treated from the same image in *DS*_1_. From example, the three images of digit 5 in *DS*_1_, *DS*_2_ and *DS*_3_ in the upper panels of Figure S1a are ‘corresponding’, because the images in *DS*_2_ and *DS*_3_ are treated (i.e, blurred and sharpened respectively) from the same image in *DS*_1_. (b) The training scheme of Figure 4e-g and Figure S2c-e. We used the same dataset (*DS*_1_, *DS*_2_ or *DS*_3_) as the input to the VAE encoder and the target of the output of the VAE decoder. (c-e) The quality of VAE-generated images, similar to Figure 4e-g. Compared this figure with Figure 4e-g and Figure S2c-e, we notice that the quality of the generated sketches depends on the dataset used as the target of the output of the VAE decoder, but hardly depends on the dataset used as the input to the VAE encoder.

**Figure S5:**
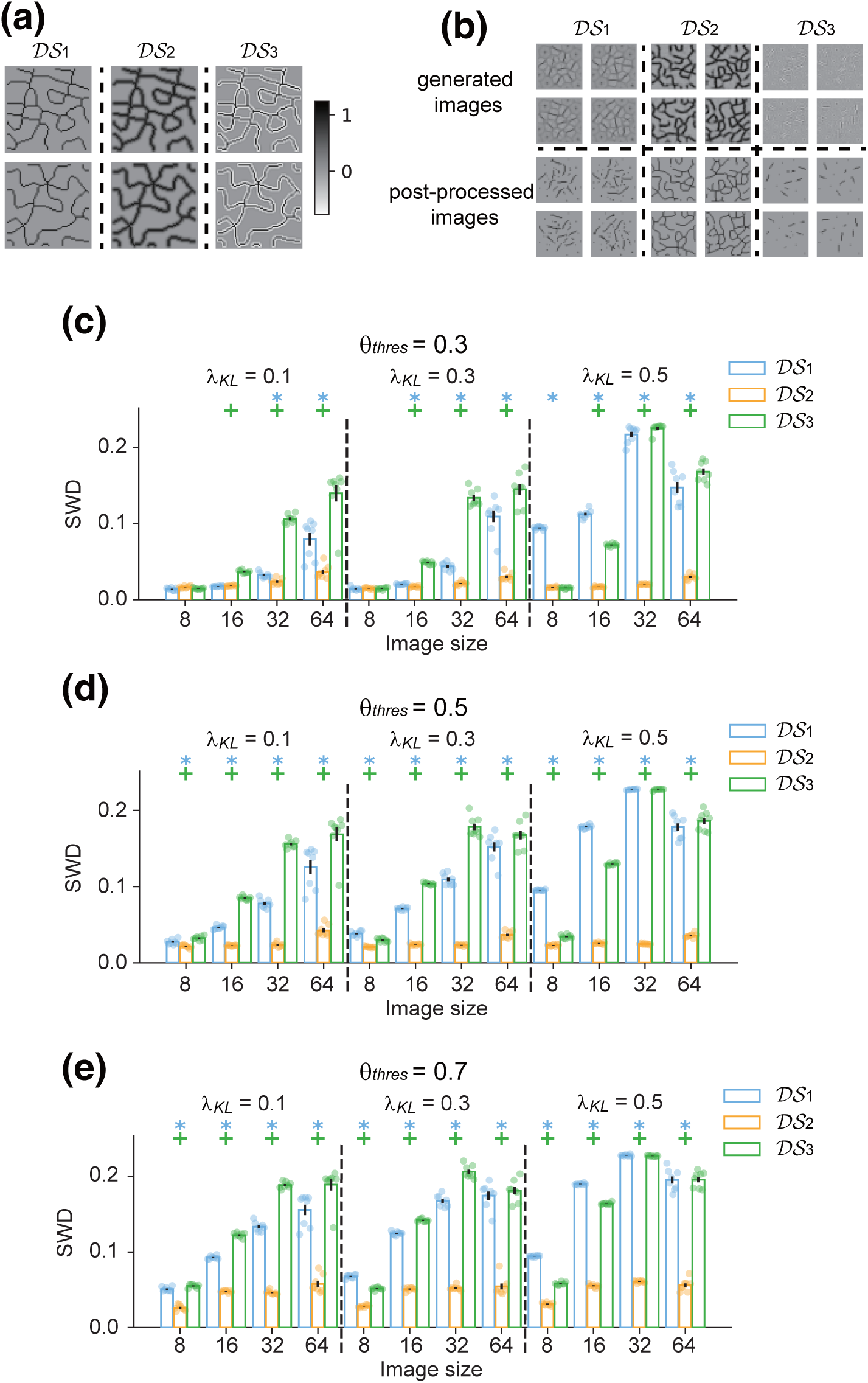
Generating thin-line textures. (**a**) Examples of *DS*_1_, *DS*_2_ and *DS*_3_ treated from OpenSimplex2 textures. *DS*_1_ and *DS*_2_ have been illustrated in Figure 4i of the main text. Image size is 64 *×* 64. (**b**) Examples of VAE-generated images (upper row) and post-processed images (lower row). Note that *DS*_3_ results in the best quality of the post-processed images. (**c-e**) SWD for different image sizes at different *λ_KL_* values, when *θ_thres_* = 0.3 (panel **c**), 0.5 (panel **d**) or 0.7 (panel **e**). Errorbars representing s.e.m. over 8 VAE configurations. Blue asterisk (or green plus sign) means that SWD for *DS*_2_ is significantly (*p <* 0.05) smaller than that for *DS*_1_ (or *DS*_3_) at the same patch size and *λ_KL_*.

**Figure S6:**
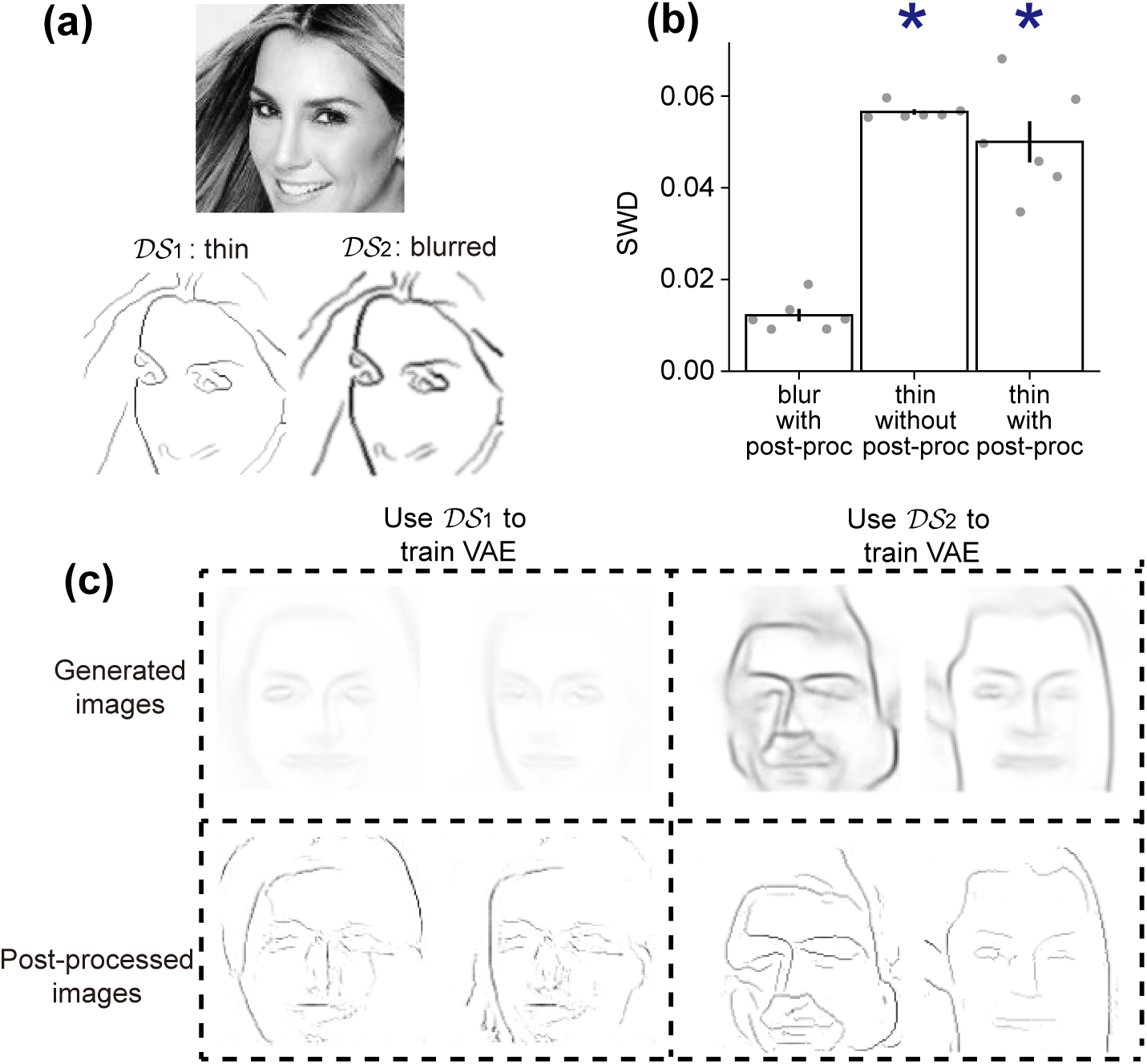
Generating continuous-valued thin contours of CelebA images using VAE. (**a**) A gray-scale CelebA image (top), its thin contours (*DS*_1_, bottom left) and its blurred contours (*DS*_2_, bottom right). (**b**) Sliced Wasserstein distance (SWD) between the resulted images and the images in *DS*_1_. Left bar: We train the VAE using blurred contours and post-process the VAE-generated images to infer the thin lines. Middle bar: We train the VAE using thin contours and do not post-process the VAE-generated images. Right bar: We train the VAE using thin contours and post-process the VAE-generated images. (**c**) Examples of the VAE-generated (upper row) and post-processed (lower row) images, using *DS*_1_ (left column) or *DS*_2_ (right column) to train the VAE.

### 3 Generating continuous-valued contours of CelebA faces using VAE

In **Figure 5**, the CelebA contours used to train the VAE are constant-valued: all pixels on the thin lines have the same intensity (see *DS*_1_ in **Figure 5a**). Here we trained VAE to generate continuous-valued CelebA contours in which different pixels on the thin line have different intensities. We found similar results to the constant-valued case: the best generation quality is realized by training VAEs using blurred contours and inferring thin contours from the VAE-generated images, instead of training VAEs using thin contours directly (Figure S6).

#### Preparing the contour dataset

The continuous-valued contours *C*_continuous_ of a CelebA face *I* was defined as

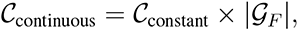

where *C*_binary_ is the constant-valued contour obtained in the same way as *DS*_1_ of **Figure 5a** (see Methods), and *|G_F_ |* is the magnitude of the intensity gradients of the corresponding CelebA image *I*, calculated using the ‘Sobel’ routine of OpenCV.

#### The network structure of VAE

The structure of the VAE to generate continuous-valued contours was the same as the structure of the VAE we used for constant-valued contours (**Figure 5**), see Methods . *λ_KL_* = 0.5.

#### The post-processor

Continuous-valued thin contours are inferred from the VAE-generated images using a post-processor. Similar to the V1 model (**Figure 2f, blue box**, see Methods for details), this post-processor was a feedforward neural network with U-Net structure. It had a similar structure to the VAE, except that there was a direct link between the *i*th (counting from the input side) layer of the encoder and the *i*th (counting from the output side) layer of the decoder. The post-processor was also trained by mean-square error plus an adversarial loss, see Methods .

### 4 Generating constant-valued contours of CelebA faces using GAN

To further examine the generalizability of our findings, we trained generative adversarial networks (GAN), another dominating image-generation paradigm, to generate constant-valued CelebA contours. We found a similar result: the best generation quality is realized by training GANs using blurred contours and inferring thin contours from the GAN-generated images, instead of training GANs using thin contours directly (Figure S7).

**Figure S7:**
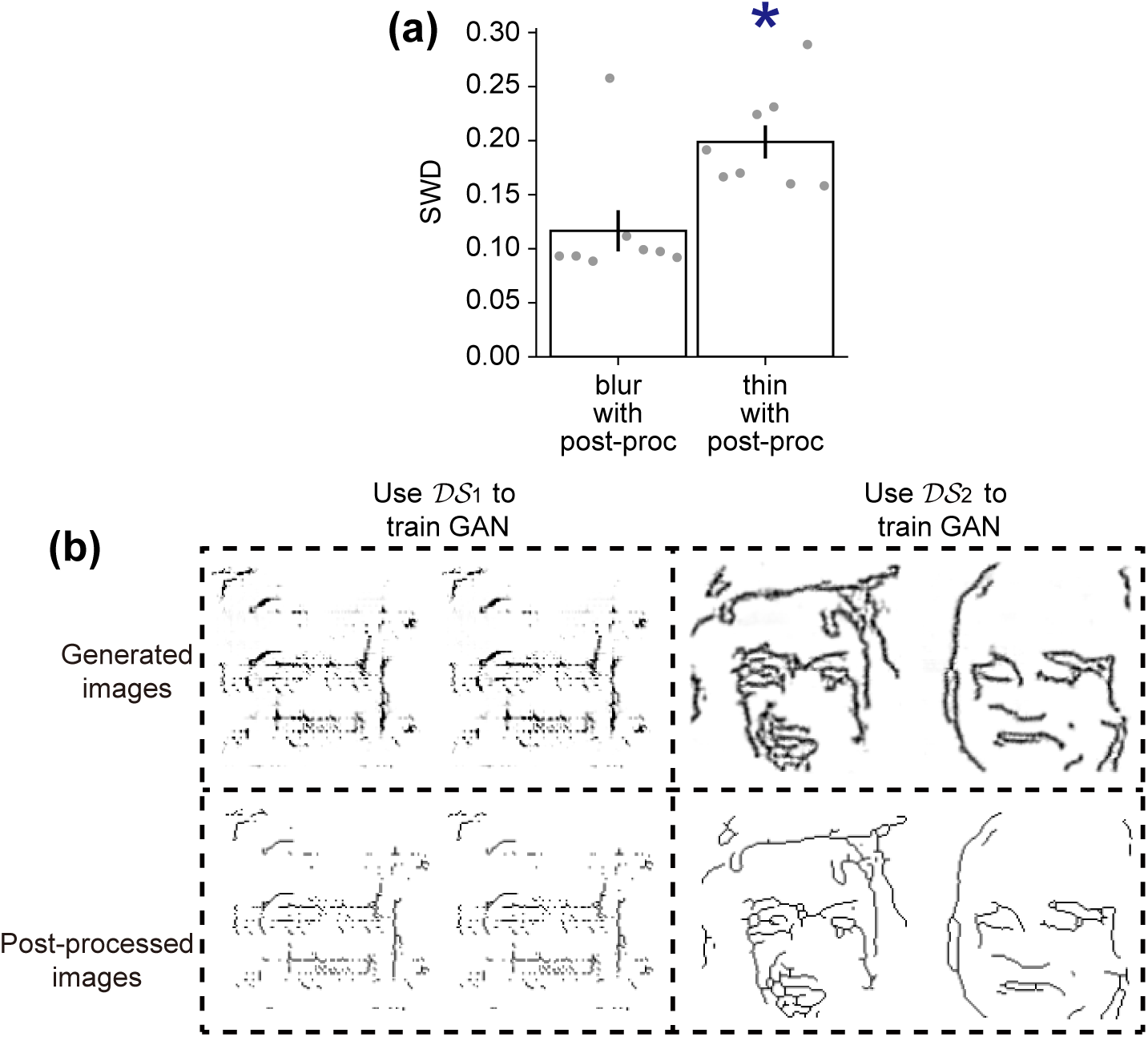
Generating constant-valued thin contours of CelebA images using GAN. (**a**) Sliced Wasserstein distance (SWD) between the resulted images and the images in *DS*_1_, if we train the GAN using blurred (left bar) or thin (right bar) contours and post-process the GAN-generated images to infer the thin lines. Error bars represent s.e.m. over 8 GAN configurations. (**b**) Examples of the GAN-generated (upper row) and post-processed (lower row) images, using *DS*_1_ (left column) or *DS*_2_ (right column) to train the GAN.

#### Preparing the contour dataset

We used constant-valued contours of CelebA images to train GANs. These contours were obtained in the same way as **Figure 5**, see Methods for detail.

#### The structure of GAN

The generative network was a feedforward network. Its input had spatial size 1 *×* 1 and 128 channels; its output had spatial size 128 *×* 128 and 1 channel. It had 6 hidden layers with spatial sizes 2 *×* 2, 4 *×* 4, 8 *×* 8, 16 *×* 16, 32 *×* 32 and 64 *×* 64 and channel numbers 64, 128, 256, 512, 512 and 512 respectively. Adjacent layers were connected by transposed convolutional kernels, followed by batch normalization and leaky ReLU activation function.

The adversarial network was also a feedforward network. Its input had spatial size 128 *×* 128 and 1 channel, its output had spatial size 1 *×* 1 and 1 channel. It had 5 hidden layers with spatial sizes 64 *×* 64, 32 *×* 32, 16 *×* 16, 8 *×* 8 and 4 *×* 4 and channel numbers 16, 32, 64, 128 and 256 respectively. Adjacent layers were connected by transposed convolutional kernels, followed by batch normalization and leaky ReLU activation function. The output was treated by the sigmoid activation function.

The parameters in the generative network were updated by minimizing

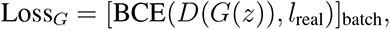

where *G*(*z*) is the output of the generative network after receiving a Gaussian noise *z*, *D*(*·*) is the output of the adversarial network, *l*_real_ = 1 is the label for real images, BCE(*x, y*) means the binary cross entropy between *x* and *y*, [*·*]_batch_ means average over batches.

The parameters in the adversarial network were updated by minimizing

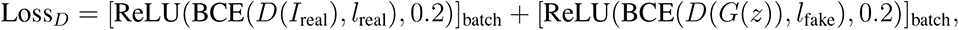

where *I*_real_ indicates a real image, *l*_fake_ = 0 is the label for fake images, ReLU(*x,* 0.2) equals *x* if *x >* 0.2 but equals zero otherwise. We introduced this ReLU function here to stop training the adversarial network if the adversarial network is already powerful enough, which improves the stability of GAN.

#### Post-processing the generated images

The images generated by GAN were post-processed similarly to **Figure 4c**, with the binarization threshold *θ_thres_* = 0.35.

